# Structural insights into crista junction formation by the Mic60-Mic19 complex

**DOI:** 10.1101/2022.03.30.486340

**Authors:** Tobias Bock-Bierbaum, Kathrin Funck, Florian Wollweber, Elisa Lisicki, Janina Laborenz, Jeffrey K. Noel, Manuel Hessenberger, Alexander von der Malsburg, Karina von der Malsburg, Carola Bernert, Séverine Kunz, Dietmar Riedel, Hauke Lilie, Stefan Jakobs, Martin van der Laan, Oliver Daumke

**Author notes:** Corresponding author. (OD); (MvdL). these authors contributed equally (Tobias Bock-Bierbaum, Kathrin Funck).

## Abstract

Mitochondrial cristae membranes are the oxidative phosphorylation sites in cells. Crista junctions (CJs) form the highly curved neck regions of cristae and are thought to function as selective entry gates into the cristae space. Little is known about how CJs are generated and maintained. We show that the central coiled-coil domain of the mitochondrial contact and cristae organizing system (MICOS) subunit Mic60 forms an elongated, bow tie-shaped tetrameric assembly. Mic19 promotes Mic60 tetramerization via a conserved interface between the Mic60 mitofilin and Mic19 CHCH domains. Dimerization of mitofilin domains exposes a crescent-shaped membrane-binding site with convex curvature tailored to interact with curved CJ necks. Our study suggests that the Mic60-Mic19 subcomplex transverses CJs as a molecular strut, thereby controlling CJ architecture and function.

## Main Text

Mitochondria are highly dynamic double membrane-bound organelles crucial for cellular metabolism, energy conversion, signaling and apoptosis (*1–8*). They are characterized by extended and intricately folded inner membrane structures termed cristae that were described in the early days of electron microscopy and later recognized as the main sites of oxidative phosphorylation. Cristae are highly adaptive and variable in shape and size depending on cell type, metabolic state and developmental stage (*3, 7, 9, 10*). Key determinants for cristae morphology are oligomeric F_1_F_o_-ATP synthase complexes that shape the tips and rims of cristae (*11*), whereas filaments of dynamin-like Mgm1/OPA1 are thought to stabilize and deform cristae from the intracristal space in an energy-dependent manner (*12, 13*). Cristae are connected to the mitochondrial envelope via crista junctions (CJs) (*9, 14–18*) (fig. S1). These highly curved tubular openings with a circular or slit-like cross-section have been suggested to function as selective pores for proteins and metabolites controlling passageways in and out of the intracristal space (*14, 15, 19*).

The conserved multi-subunit mitochondrial contact site and cristae organizing system (MICOS) localizes to CJs (*20–25*). It is crucial for the formation and stabilization of CJs from yeast to humans and likely plays an important role for the regulation of CJ permeability. MICOS is composed of the Mic60 and Mic10 subcomplexes that both possess membrane shaping activity (fig. S1) (*26–32*). Mic60 is anchored in the IMM via an N-terminal transmembrane (TM) segment and exposes a large domain into the intermembrane space (Fig. 1A) that associates with Mic19 (and in metazoa additionally Mic25). The Mic60 module of MICOS links CJs to the mitochondrial outer membrane through formation of membrane contact sites with different partner protein complexes, like the sorting and assembly machinery for β-barrel proteins (SAM complex) (*9, 10, 21-23, 33-37*). Mic12 in yeast or MIC13/QIL1 in higher eukaryotes connect the Mic60 and Mic10 modules of MICOS (*28, 38, 39*). Loss of MICOS components leads to a massively altered cristae morphology in all organisms examined so far. The strongest phenotypes with a nearly complete loss of CJs and accumulation of detached sheets of lamellar cristae membranes are observed upon ablation of Mic60 and Mic10 (*9, 17, 20-23, 25, 40*). How exactly MICOS controls CJ architecture and function is, however, unclear since no structural information of any MICOS component is available.

**Fig. 1.**
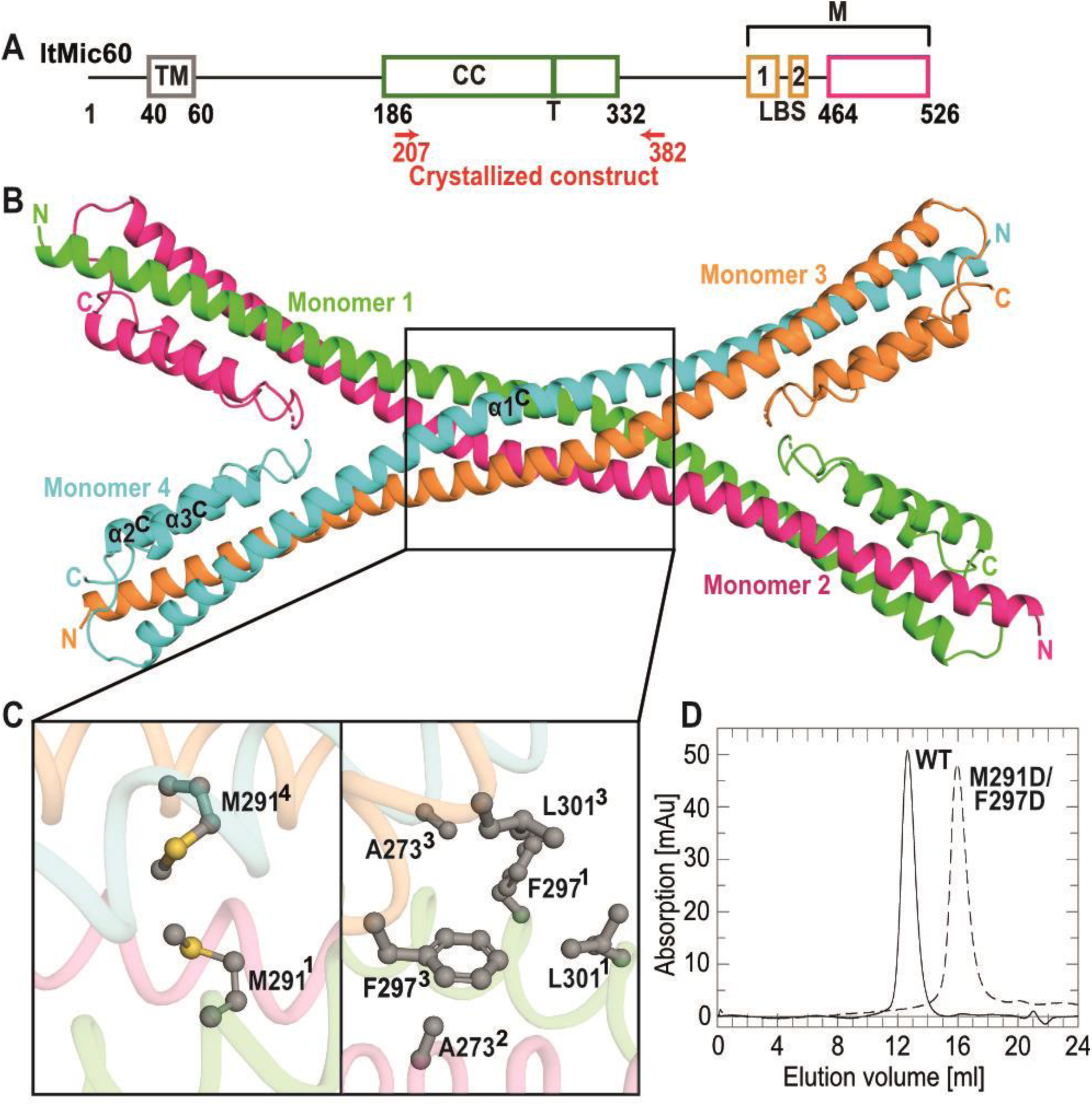
Mic60 coiled-coil domain forms an antiparallel tetramer. (**A**) Domain architecture of Mic60 (amino acid numbers refer to Mic60 from *Lachancea thermotolerans*). TM: transmembrane helix, CC: coiled-coil, LBS 1+2: lipid binding site, M: mitofilin domain. The green bar indicates the tetramer interface (T) and the magenta box denotes the previously reported, sequence-based boundaries of the mitofilin domain. (**B)** Cartoon representation of the Mic60 coiled-coil domain (ltMic60_CC_). N- and C-termini of each monomer are labelled. (**C)** Close up of the tetramer interface. (**D)** SEC profile of ltMic60_CC_ and ltMic60_CC_^M291D/F297D^

Here, we found that the central coiled coil domain of Mic60 forms a bow-tie shaped tetrameric assembly. Mic19 promotes Mic60 tetramerization. The C-terminal mitofilin domains of Mic60 dimerize to form two crescent-shaped membrane-binding modules on each side of the coiled-coil. Our structural study suggests that the Mic60-Mic19 complex traverses CJs, therefore controlling their diameter and function.

## Results

### The coiled-coil domain of Mic60 forms an antiparallel tetramer

In tomograms of fixed *Saccharomyces cerevisiae* (*S. cerevisiae*), we frequently observed filamentous density in the mitochondrial CJs (fig. S1c). We reasoned that this density may constitute part of the MICOS complex and, due to its elongated shape, the predicted coiled-coil (CC) region of Mic60. To obtain structural information, we determined the crystal structure of the Mic60 coiled-coil domain from the thermostable yeast *Lachancea thermotolerans* (*L. thermotolerans*, amino acids 207 to 382; ltMic60_CC_) (Fig. 1A, table S1 and S2). LtMic60_CC_ forms an elongated α-helix (α1C) with two short α-helices (α2C and α3C) tightly packed onto the C-terminal ends of α1C (Fig. 1B). In agreement with size exclusion chromatography and Blue Native PAGE (BN-PAGE) analysis (fig. S2A), four ltMic60_CC_ molecules assemble into a tetramer via a hydrophobic, highly conserved interface (Fig. 1B, C and fig. S3A, B and 4). An antiparallel dimeric coiled-coil further dimerizes to form a bow tie-shaped tetrameric assembly. In agreement with the structure, a double amino acid substitution in this interface (M291D/F297D) leads to disruption of the ltMic60_CC_ tetramer into monomers (Fig. 1D, fig. S2A).

### Mic19 promotes Mic60 tetramerization to stabilize crista junctions

Since longer constructs of ltMic60 could not be expressed in a soluble form, we resorted to Mic60 from *Chaetomium thermophilum* (*C. thermophilum*, ct) for further biochemical analysis. In agreement with previous data (*30*), an almost full-length construct of ctMic60 excluding the TM region (residues 208-691, ctMic60_sol_) migrates as a dimer in BN-PAGE, with some minor higher order assemblies (Fig. 2A, B and table S2). Strikingly, addition of purified *C. thermophilum* Mic19 induced the formation of a heteromeric species, likely containing four molecules of Mic60 and Mic19 each (Fig. 2B).

**Fig. 2.**
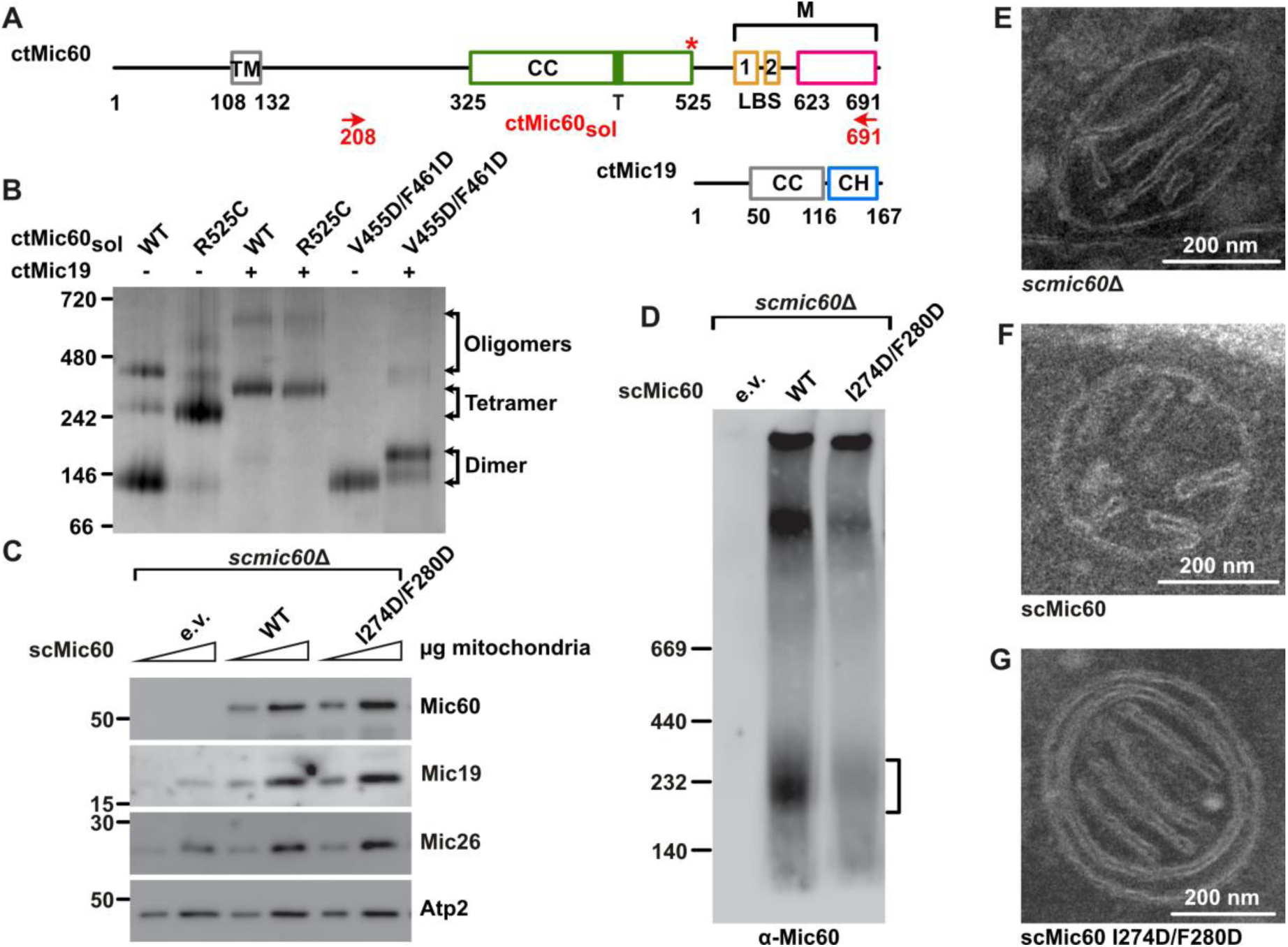
Mic19 promotes Mic60 tetramerization to stabilize crista junctions. (**A**) Domain architecture of ctMic60 and ctMic19, as in Fig. 1A. CH: coiled-coil-helix-coiled-coil-helix domain. The red asterisk indicates the position of the artificially introduced disulfide bridge. **(B)** Representative BN-PAGE analysis showing oligomerization of different ctMic60_sol_ variants and their complexes with ctMic19. **(C)** Steady state levels of selected mitochondrial proteins in the indicated *S. cerevisiae* strains determined by SDS-PAGE and Western Blot; e.v. empty vector. **(D)** BN-PAGE and immunoblot analysis of Mic19-Protein A immunoprecipitation elution fractions showing the oligomeric state of isolated Mic60-containing complexes. The deduced position of the tetrameric Mic60 species is indicated by the bracket. **(E-G)** Ultrathin sections of *S. cerevisiae* mitochondria in the indicated strains (see Fig. 4D and fig. S7E for quantification).

To prove the involvement of the tetrameric interface in this assembly, we introduced a structure-based disulfide bridge in ctMic60_sol_ (R525C) which can only form in the tetrameric context (fig.S3C). Indeed, under oxidizing conditions, ctMic60_sol_^R525C^ formed a tetramer even in the absence of Mic19 (Fig. 2B, fig. S2B). In the presence of Mic19, the assembly was shifted to a hetero-oligomeric complex of comparable size to ctMic60_sol_-Mic19 in BN-PAGE. This indicates that the cross-link stabilizes a native form of the Mic60-Mic19 assembly. Furthermore, a double amino acid substitution in the tetrameric interface, V455D/F461D, greatly reduced higher order assembly of Mic60 in the absence and presence of Mic19 (Fig. 2B).

CtMic60_sol_ co-sedimented with Folch liposomes derived from bovine brain lipids and dragged ctMic19 into the pellet fraction (fig. S2C, D) (*30*). CtMic60_so_ ^R525C^ and ctMic60_so_ ^V455D/F461D^ proteins co-sedimented with liposomes to a similar extent as ctMic60_sol_. Interestingly, the ctMic60_sol_^V455D/F461D^ variant showed reduced recruitment of Mic19 to Folch liposomes (fig. S2D) and a 30-fold reduced affinity to Mic19 in isothermal titration calorimetry (ITC) experiments compared to ctMic60_sol_ (fig. S6A, B, L). This suggests that oligomerization of Mic60 via the tetrameric interface is required for a tight interaction with Mic19.

To analyze the physiological role of Mic60 tetramerization, we employed *S. cerevisiae* (sc) as a model. Mic60-deficient *S. cerevisiae* cells showed massively reduced levels of Mic19 (Fig. 2C), as previously described (*21, 23*). Expression of a tetramer-disruptive scMic60 variant (I274D/F280D, fig. S4) restored mitochondrial accumulation of Mic19 in these cells (Fig. 2C), but interfered with the appearance of Mic60-containing tetrameric complexes, as revealed by BN-PAGE (Fig. 2D). Mic60-deficient cells showed the expected loss of CJs, which was rescued by re-expression of scMic60 (Fig. 2E, F, fig. S7E, F). Strikingly, re-expression of tetramerization-defective scMic60 variant in *mic60Δ* cells did not restore CJ architecture (Fig. 2G). In EM tomograms of these mitochondria, the few remaining CJs did not show a filamentous density (fig. S1D), suggesting a role of the Mic60 tetramer in the formation of this structure. We conclude that tetramerization of Mic60 is required for proper CJ formation.

### The Mic60 mitofilin domain binds to the Mic19 CHCH domain via a conserved interface

We next aimed to characterize the molecular basis of the Mic60-Mic19 interaction, which requires the C-terminal mitofilin domain of Mic60 and the C-terminal CHCH (coiled-coil-helix-coiled-coil-helix) domain of Mic19 (*30*). Because isolated mitofilin domain constructs tended to precipitate after purification, we determined the crystal structure of a fusion construct (termed Mito1_CHCH), containing the C-terminal region of the ctMic60 mitofilin domain linked to the ctMic19 CHCH domain (table S1 and S2).

Each Mito1_CHCH monomer consists of one α-helix from the mitofilin domain (α3M) and α1CH and α2CH of the Mic19 CHCH domain, which together form a three helical bundle (Fig. 3A). The mitofilin-CHCH domain interface has an area of 660 Å^2^ and is dominated by conserved, hydrophobic interactions (Fig. 3B, fig. S3E, S4 and S5). Mutagenesis of the hydrophobic residues in the context of the longer ctMic60_sol_ and ctMic19 construct impaired formation of higher-order oligomers in BN-PAGE (fig. S2E) and strongly reduced binding affinity in ITC experiments (see color code in Fig. 3B, fig. S6A, E-M) as well as Mic60-mediated recruitment of Mic19 to liposomes (Fig. 3C, fig. S2D). Thus, the interaction in the fusion construct faithfully reflects the interaction of Mic60 and Mic19. In contrast, disruption of the peripheral polar interactions in the interface showed only minor effects on oligomerization and a moderate reduction in binding affinity (Fig. 3B, fig. S2E and S6).

**Fig. 3.**
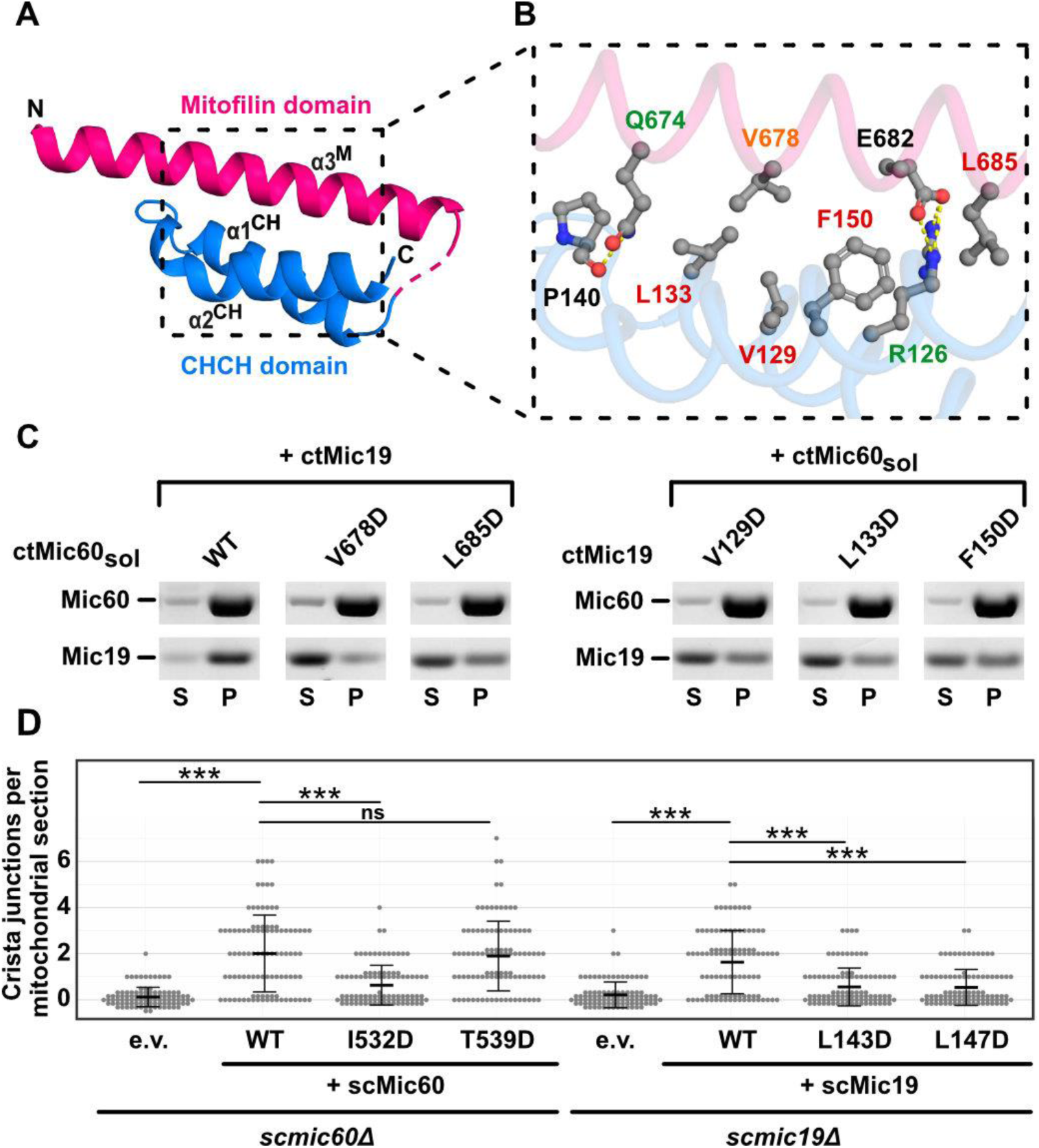
Structural characterization of the Mic60 and Mic19 interaction. **(A)** Structure of Mito1_CHCH fusion construct. Domains are colored as in Fig. 2A. The unresolved linker region is indicated by a dotted line. **(B)** Magnification of the ctMic60-ctMic19 interface. Label colors represent the effect of single amino acid substitutions, as analyzed by ITC experiments: Green - up to 15x reduced; orange - up to 100x reduced, red - no binding, compared to ctMic60_sol_ + ctMic19; see fig. S6 for ITC data. **(C)** Liposome co-sedimentation assays of ctMic60_sol_ and ctMic19. S, supernatant; P, pellet (see also fig. 2C, D). **(D)** Quantification of crista junctions per mitochondrial section in *S. cerevisiae* mitochondria. *p ≤ 0.05; **p ≤ 0.01, ***p ≤ 0.001. ns: not significant, see also fig. S7F.

When re-introduced into Mic60-deficient *S. cerevisiae* cells, the scMic60 I532D variant with a defective Mic60-Mic19 interface showed reduced levels of mitochondrial Mic19, in comparison to cells re-expressing wild-type scMic60 (fig. S4, S7A). Isolated Mic60-containing oligomeric complexes were reduced in this mutant (fig. S7D). Accordingly, scMic60 I532D-containing mitochondria were almost devoid of CJs (Fig. 3D, Fig. S7F). By contrast, the scMic60 T539D substitution at the periphery of the interface did not induce these effects.

*S. cerevisiae* Mic19 variants with amino acid substitutions in the Mic60-Mic19 interface (scMic19 L143D and L147D, fig. S5) showed reduced accumulation of Mic19 in mitochondria and also hardly any CJs, when re-expressed in a *MIC19* deletion strain (Fig. 3D, fig. S7B, F). These data reveal the critical importance of the hydrophobic Mic60-Mic19 interface for protein stability and MICOS integrity in living cells.

### The mitofilin dimer forms a convex membrane binding site

Besides the N-terminal TM anchor, Mic60 interacts with membranes via two distinct lipid binding sites (LBS 1+2) in the C-terminal region of the protein (Fig. 1A, 2A) (*30*). Constructs including LBS 1+2 did not crystallize. Co-evolution analysis (*41*) predicted that LBS 2 is flexible (fig. S3D) and might therefore interfere with crystallization. Sourcing this information, we determined the crystal structure of the mitofilin domain including LBS 1, but without LBS 2, again fused to the CHCH domain of ctMic19 (Mito2_CHCH, table S1 and S2).

The Mito2_CHCH structure revealed that the mitofilin domain is built of a four-helix bundle: α1M, α2M and the LBS 1 from one monomer interact with α3M from an opposing monomer to form an inter-domain swapped dimer with an interface area of 2,000 Å2 (Fig. 4A). The interaction of α3M with the CHCH domain of Mic19 is identical to the previously described structure containing the truncated mitofilin domain construct (Fig. 3A). L676 in the dimer interface points into a hydrophobic pocket of the interacting monomer (Fig. 4A). In agreement with the structural data, the Mito2_CHCH construct was a dimer in analytical ultracentrifugation (AUC) experiments, whereas the L676D amino acid substitution rendered the protein monomeric (Fig. 4B). In the longer ctMic60_sol_ construct (amino acids 208-691), the L676D variant showed a similar behavior as the wily-type protein in BN-PAGE, forming mostly a dimer (Fig. 4C). Liposome binding of the variant was also comparable to that of unmodified ctMic60_sol_ (fig. S2C). However, complex formation with ctMic19 in BN-PAGE (Fig. 4C) and ITC (fig. S6C, L), and Mic60-mediated Mic19 recruitment to liposomes was reduced (fig. S2D), indicating that dimerization of the Mic60 mitofilin domain supports Mic19 recruitment. Simultaneous disruption of the tetrameric and dimeric interface in the V455D/F461D/L676D variant had an even more drastic effect, completely preventing higher-order oligomer formation of ctMic60_sol_ alone and in complex with ctMic19 (Fig. 4C. In addition, Mic60-dependent Mic19 recruitment to liposomes was severely affected (fig. S2D). Having both interfaces disrupted, the number of CJs per mitochondrial section in the respective *S. cerevisiae* variant (scMic60_I274D/F280D/V530D_) were equally reduced as in the complete *MIC60* knockout strain (Fig. 4D, fig. S7C, F), showing the additive effect of both assembly sites for tetramerization.

**Fig. 4.**
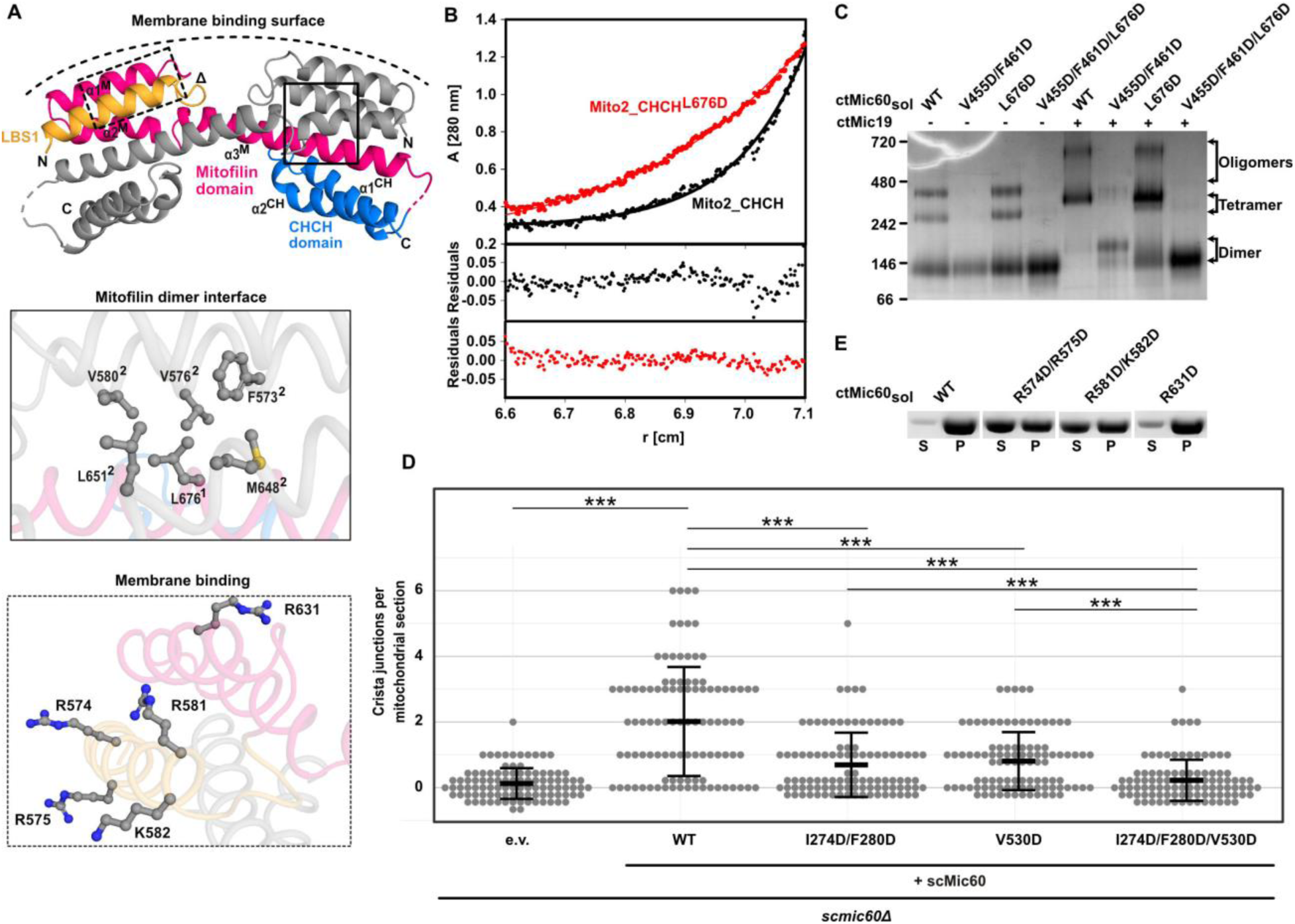
The mitofilin dimer forms a convex membrane binding site. **(A)** Cartoon representation of the mitofilin dimer in complex with the Mic19 CHCH domain (Mito2_CHCH). One monomer is colored as in Fig. 2A. Dimer interface residues (boxed in straight lines) and potential membrane binding residues (boxed in dotted lines) are shown in the close-up views. The deduced curved membrane binding surface is indicated. **(B)** Association states of Mito2_CHCH (black) and Mito2_CHCH^L676D^ (red) were analyzed at a concentration of 1 mg/ml by sedimentation equilibrium analysis. Upper panel shows the original data (dots) and the fit of the data (line): Mito2_CHCH: molecular mass (Mr) = 34 ± 4 kDa. Mito2_CHCH^L676D^: Mr = 16 ± 2 kDa (molecular weight of the monomer: 17 kDa). The lower panel shows the deviation of the fit to the data. **(C)** BN-PAGE analysis of different ctMic60_sol_ variants and their complexes with ctMic19. **(D)** Quantification of crista junctions per mitochondrial section in *S. cerevisiae* mitochondria. *p ≤ 0.05; **p ≤ 0.01, ***p ≤ 0.001. ns: not significant, see also (see fig. S7F). **(E)** Liposome co-sedimentation assays of ctMic60_sol_ and membrane binding variants. S: supernatant, P: pellet (see also fig. S2C).

The dimeric arrangement positions the two positively charged LBS 1-helices on the outside of the dimer on a convex membrane binding surface (Fig. 4A). Replacement of the positively charged amino acid residues on this convex surface led to reduced membrane binding (Fig. 4A, E; table S2 and fig. S2C), supporting the idea that the convex surface in the mitofilin dimer comprises the membrane binding site of Mic60.

## Discussion

Our study suggests a structural model of how the Mic60-Mic19 subcomplex governs CJ formation and function (Fig. 5A, B). Each of the widely separated ends of the antiparallel tetrameric coiled-coil of Mic60 harbors two C-termini. Since the connection to the mitofilin domain is short, the two dimeric membrane-binding sites of the mitofilin domain must be localized on opposite sides of the coiled-coil. In a cellular context, this implies that the tetrameric coiled-coil spans over the CJ, and the two mitofilin domain dimers bind to opposite membrane surfaces in the CJs (Fig. 5A, B). The convex-shaped membrane-binding sites of the mitofilin dimer would be complementary to the membrane curvature of the CJs. The N-terminal TM regions of Mic60 further anchor the complex into the CJ membrane, whereas the N-terminal region of Mic19 together with parts of Mic60 could reach over to the OMM and, by interacting with SAM and TOM complexes, form membrane contact sites (fig. S1). Importantly, our model explains the uniform diameters of circular or slit-like CJs, which would be governed by the length of the traversing tetrameric coiled-coil of the Mic60-Mic19 complex. By spanning across CJs, the Mic60-Mic19 complex is tailored to serve as a physical barrier preventing the free diffusion of proteins in and out of the cristae space. In fact, super-resolution microscopy of human Mic60 suggests that up to ten Mic60 molecules are located at one CJ (*42*), further supporting the idea of an arch dome-like assembly that vaults the entry into cristae. Of note, an intrinsically disordered region is predicted between the N-terminal TM and the central coiled-coil domain of Mic60 proteins (Fig. 5A, B) (*43, 44*). Similar to the FG repeats in nuclear pore complexes, this region may contribute to the formation of a sieve-like diffusion barrier (*45*) to control selective passage through CJs.

**Fig. 5.**
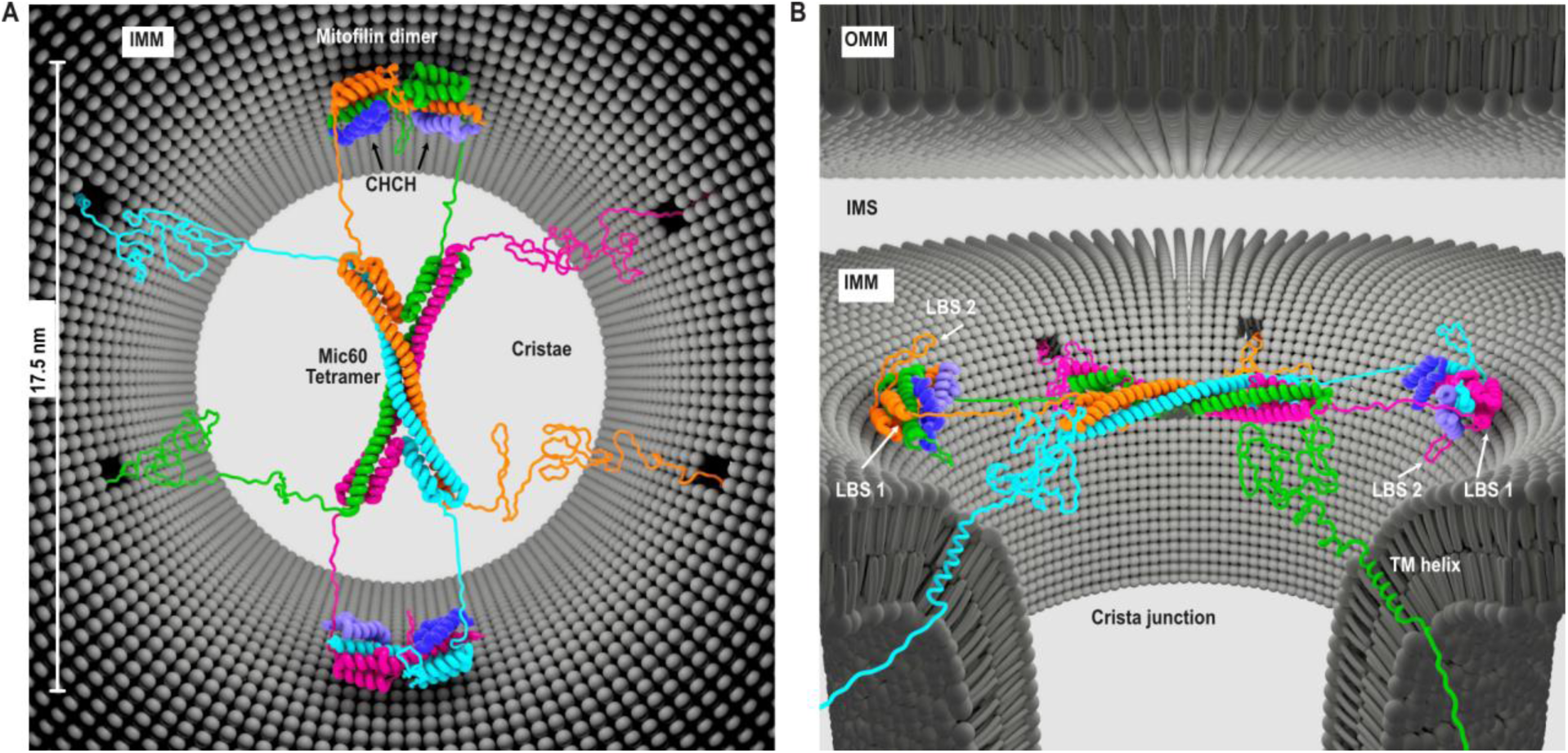
Model of Mic60-Mic19 function at CJs. **(A)** Top view and **(B)** side view showing the proposed architecture of the Mic60-Mic19 complex at CJs. Each monomer has a different color. Regions not determined by X-ray crystallography are modelled as unstructured elements.

Our model rationalizes how the Mic60-Mic19 complex governs the formation of CJ architecture and suggests how it may control protein and metabolite diffusion by acting as a tetrameric strut spanning across a CJ.

## Acknowledgments

We thank Dr. Yvette Roske for help with crystallographic data collection, Dr. Erik Werner from Research Network Services Ltd., Berlin, Germany, for the creation of the model figure, and the entire BESSY team for generous support during data collection at beamlines MX14.1, MX14.2 or MX14.3. We thank Christina Schiel for preparing ultrathin sections of *S. cerevisiae* cells (Electron microscopy, MDC), Dr. Anja Schütz (Protein Production & Characterization Platform, MDC) for support in the mass spectrometry analyses, Sibylle Jungbluth (University of Homburg) for assistance with genetic manipulation of *S. cerevisiae* and biochemical analysis of mitochondria and Stefan Stoldt (Max Planck Institute For Biophysical Chemistry, Göttingen) for the preparation of *S. cerevisiae* for electron tomography.

## Author contributions

TBB and KF designed MICOS constructs, grew crystals, solved their structures and performed biochemical experiments, with support from CB, EL. FW, JL, AvdM and KvdM performed structure-based functional experiments in *S cerevisiae*. JN conducted and analyzed evolutionary coupling predictions, EL, SK and DR provided EM analyses of *S cerevisiae* mitochondria. MH designed the Mic60-Mic19 fusion construct and grew initial crystals. HL performed AUC analyses. TBB, KF, FW, SJ, MvdL, and OD designed research and interpreted structural, biochemical and EM data. TBB, KF, MvdL and OD wrote the manuscript with inputs from all authors

## Data and materials availability

All data are available in the main text or the supplementary materials. The atomic coordinates of ltMic60CC, Mito1_CHCH and Mito2_CHCH have been deposited in the Protein Data Bank with accession numbers 7PUZ, 7PV0, 7PV1.

## Supplementary Materials

### Materials and Methods

#### Cloning and plasmids

To obtain structural information, about 200 Mic60 and Mic19 constructs from different species were cloned and expressed. Codon-optimized constructs of *L. thermotolerans* Mic60 (ltMic60; UniProt ID: C5E325, synthesized by Eurofins Genomics), *C. thermophilum* Mic60 (ctMic60; UniProt ID: G0SHY5) and Mic19 (ctMic19; UniProt ID: G0S140) were cloned into the pET26b vector (ltMic60) or modified pET28a (ctMic60 and ctMic19), encoding a C-terminal His6- tagged and a HRV-3C protease cleavable N-terminal His6-tagged fusion construct, respectively. Constructs Mito1_CHCH and Mito2_CHCH were generated using overlap extension PCR and cloned into pETDuet™-1 vector (Merck) encoding a HRV-3C protease cleavable N-terminal His6- tagged fusion protein (*48*). Variants of ltMic60, ctMic60 and ctMic19 were generated using site directed mutagenesis (*49*).

#### Expression and Purification

Expression plasmids were freshly transformed into chemical competent *E. coli* BL21 (DE3) cells. Protein expression was carried out in terrific broth supplemented with 50 µg/ml kanamycin or 100 µg/ml ampicillin. The cultures were grown until OD600 reached 0.8 at 37 °C and 80 rpm, protein expression subsequently induced by the addition of 300 µM isopropyl β-D-1-thiogalactopyranoside (IPTG) and incubated at 20 °C for another 20 h. The cells were centrifuged at 4,000 *g*, collected and frozen at -20 °C until needed.

LtMic60 expression plasmid containing cells were diluted in lysis buffer (50 mM HEPES/NaOH pH 8.0; 500 mM NaCl, 20 mM imidazole, 10% glycerol, 1 mM 3,3′,3′′-phosphanetriyltripropanoic acid (TCEP), and 1 mg/ml DNaseI (Roche)) prior to disruption using a microfluidizer (Microfluidics). To remove insoluble parts, the solution was centrifuged at 100,000 *g*, 4 °C and 45 min. The cleared supernatant was applied onto a prepacked Ni^2+^ sepharose High Performance IMAC resin (GE Healthcare Life Science) containing gravity flow column loaded with 100 mM nickel sulfate and equilibrated in lysis buffer. The column was washed using lysis buffer and bound proteins eluted with lysis buffer containing 50 and 500 mM imidazole, respectively. To remove the C-terminal His6-tag, Carboxypeptidase A (from bovine pancreas, Sigma) treatment was applied during overnight dialysis at 4 °C against dialysis buffer (50 mM HEPES/NaOH pH 8.0; 500 mM NaCl, and 10% glycerol). A second Ni^2+^ sepharose column was used to separate cleaved from uncleaved protein. Finally, a size exclusion chromatography using a S200 column and buffer SEC (50 mM HEPES/NaOH pH 8.0; 500 mM NaCl, 10% glycerol and 1 mM TCEP) (GE Healthcare Life Science) was applied to separate pure protein from aggregates and Carboxypeptidase A. Pure protein was concentrated to 31 mg/ml, flash frozen in liquid nitrogen and stored at -80 °C until further use.

All ctMic60 and ctMic19 constructs were purified in a similar manner as described for ltMic60. However, the lysis buffer contained 50 mM HEPES/NaOH pH 7.5; 500 mM NaCl and 20 mM imidazole. The N-terminal His_6_-tag was cleaved using recombinant His_6_-tagged HRV 3C protease during overnight dialysis using 50 mM HEPES/NaOH pH 7.5, 150 mM NaCl and 25 mM imidazole. The final size exclusion chromatography was performed using S200 and S75 size exclusion chromatography columns (GE Healthcare Life Science) equilibrated in 20 mM HEPES/NaOH pH 7.5; 150 mM NaCl for ctMic60 and ctMic19, respectively. CtMic60_sol_^R525C^ was purified in a similar manner in the presence of 2 - 5 mM DTT.

#### Mass spectrometry

Mass spectrometry analysis using liquid chromatography-electrospray ionization-quadrupole-time of flight-mass spectrometry (LC-ESI-Q-TOF-MS) indicated that a disulfide bridge was formed in all ctCHCH domain-containing constructs, as the calculated and the measured molecular mass showed a difference of exactly 2 Da. Protein intact mass analyses were conducted on an Agilent 1290 Infinity II UHPLC system coupled to an Agilent 6230B time-of-flight (TOF) LC/MS instrument equipped with an AJS (Agilent Jet Stream Technology) ion source operated in positive ion mode (denaturing conditions). Protein samples were desalted using a Zorbax 300SB-C3 guard column (2.1 × 12.5 mm, 5 μm). Protein solutions were diluted in 0.1% formic acid (in H_2_O) to approx. 0.06 mg/ml. Approx. 0.3 μg of sample was injected for each analysis. LC/MS parameters were adapted from ref. ^50^. The ion source was operated with the capillary voltage at 4000 V, nebulizer pressure at 50 psi, drying and sheath gas at 350 °C, and drying and sheath gas flow rate at 12 and 11 l/min, respectively. The instrument ion optic voltages were as follows: fragmentor 250 V, skimmer 65 V, and octopole RF 750 V. Mass spectrometry data were analyzed using the Protein Deconvolution feature of the MassHunter BioConfirm Version 10.0 software (Agilent) that uses the Maximum Entropy algorithm for accurate molecular mass calculation. Deconvolution was performed between mass range of 800 to 2,500 m/z (mass-to-charge ratio), using peaks with a ratio of signal to noise greater than 30:1. The deconvoluted mass range was set at 5 to 25 kDa and the step mass was 1 Da.

#### Analytical size exclusion chromatography

Analytical size exclusion chromatography was performed using 100 µl of a 3 mg ml^-1^ protein solution on a Superdex S200 10/300 (GE Healthcare Life Science) column at 0.5 ml/min flow rate and 4 °C. The running buffer contained 50 mM HEPES/NaOH pH 8, 500 mM NaCl.

#### Crystallization, data collection, refinement and other tools

Initial crystallization conditions were identified with the vapor diffusion method in 96 well sitting drop format at 20 °C using an automated dispensing robot (Art Robbins Instruments). Optimized and plate like protein crystals grew within 3-5 days by mixing 1 µl of 31 mg/ml ltMic60CC and 1µl reservoir containing 17.5 % PEG 1500, 0.1 M MMT buffer pH 7.1 (a 1:2:2 molar mixture of DL-malic acid, MES and Tris base) and 0.1 M D-Sorbitol. The drop was equilibrated against 500 µl of the same reservoir solution in 24 well hanging drop format at 20 °C. In order to obtain high quality diffracting protein crystals, dehydration was performed (*51*). Therefore, protein crystals were transferred into a new drop containing 35% PEG 1500, 250 mM NaCl, 0.1 M MMT buffer pH 7.2 and 0.1 D-(-)-fructose and equilibrated against 500 µl of the same solution for 24 h prior to flash cooling in liquid nitrogen.

Diffraction quality crystals of Mito1_CHCH and Mito2_CHCH grew within 2-10 days and were obtained in 96 well sitting drop format at 20 °C by automated mixing 0.2 µl protein solution (17 mg/ml - 20 mg/ml) and 0.2 µl reservoir solution (80 µl total reservoir volume). 10 min prior to crystallization trials, Mito1_CHCH was mixed with 1% trypsin [w/w] and incubated at 4 °C. The final reservoir solutions contained 33% [v/v] Jeffamine M-600, 0.1 M HEPES/NaOH pH 7.2 (Mito1_CHCH) or 30% [w/v] Jeffamine ED-2001, 0.1 M HEPES/NaOH pH 7.0 (Mito2_CHCH). After they stopped growth, crystals were directly flash cooled in liquid nitrogen. Diffraction data were collected at -173 °C and 0.9184 Å on beamline BL14.1 operated by the Helmholtz–Zentrum Berlin at the BESSY II electron storage ring (Berlin–Adlershof, Germany) (*52*) and indexed, integrated and scaled with XDSAPP (*53*). The structure of ltMic60CC and Mito1_CHCH was solved by molecular replacement using AMPLE from the CCP4 software packaging (*54, 55*). For ltMic60CC, four ideal α-helices build by 20 alanine residues have been placed and AutoBuild from the PHENIX suite was used for initial model building (*56, 57*). For Mito1 _CHCH, three ideal α-helices build by 30 alanine residues could be placed. Additional helices have been identified using Phaser-MR (*58*) from the PHENIX suite. SHELXE (*59*) and AutoBuild were used to obtain the initial model. The structure of Mito2_CHCH was solved by molecular replacement with Phaser-MR using the final refined structure of Mito1_CHCH as search model. LtMic60CC crystallized in space group P4212 with one monomer, Mito1_CHCH in P1 with six monomers and Mito2_CHCH in P21 with four monomers arranged as two dimers in the asymmetric unit. For ltMic60CC, only amino acids 235-382 are visible in the structure. Residues ctMic60^661-691^ and ctMic19^118-158^ are visible in all monomers of Mito1_CHCH. In case of Mito2_CHCH, ctMic60^565-586^, ctMic60^622-689^ and ctMic19^118-160^ are visible in all monomers.

Refinement was carried out using iterative steps of manual model building in Coot (*60*) and maximum likelihood refinement with individual B-factors, TLS and secondary structure restraints using phenix.refine (*61*). Final structure validation was carried out with MolProbity (*62*). All statistics for data collection and refinement as well as the corresponding PDB codes can be found in Extended Data Table S1.

Surface conservation plot were created using the ConSurf Server (*63*) with standard settings and multiple sequence alignments using MULTALIN64. Figures were prepared with PyMOL (The PyMOL Molecular Graphics System, Version 1.8.2.3 Schrödinger, LLC). PDBePISA web server (*65*) was used to calculate the interface area between the mitofilin domain and the CHCH domain.

#### Evolutionary coupling analysis

Structure prediction of the C-terminal region of Mic60 was done using the EVCoupling server (*41*). Structure predictions of ctMic60^557-685^ was obtained using the input sequence ctMic60^550-693^ (monomer pipeline, version 1, bitscore 0.1). The highest scoring folding candidate is shown.

#### Oxidation of cysteines

In order to artificially induce disulfide bridge formation, 30 µM of the respective protein has been dialyzed against 20 mM Tris-HCl pH 7.5 and 150 mM NaCl overnight. The next day, oxidation of free cysteines was performed using 500 µM CuSO4 for 15 min at 4 °C followed by the addition of 50 mM ethylenediaminetetraacetic acid (EDTA). The residual CuSO4 and EDTA was removed using a PD-10 column (GE Healthcare). The concentration of the final oxidized protein was set to 1 mg/ml for further analysis.

#### Blue native PAGE

Blue native PAGE (BN-PAGE) analysis of recombinant purified proteins was performed using the Native PAGE Bis-Tris system (Thermo Fisher Scientific). 5 µg (single proteins) or 5 µg (Mic60 variants) incubated for 15 min with 2 µg (Mic19 variants) were applied on 4-16% acrylamide BN-PAGE gradient gels and run for 2 h at 150 V on ice. Proteins were visualized by Coomassie Brilliant Blue (CBB) staining. Immunoprecipitated mitochondrial protein complexes eluted in digitonin buffer (see below) were mixed with 10x BN-PAGE loading buffer (5% [w/v] Coomassie brilliant blue G250, 500 mM 6-aminocaproic acid, 100 mM Bis-Tris/HCl pH 7.0) and loaded on home-made 4-13% BN-PAGE gradient gels that were run for 2-3 h at 150 V in a water-cooled Hoefer gel chamber at 6°C. Proteins were blotted on polyvinylidene difluoride (PVDF) membranes and visualized using specific antibodies together with an enhanced chemiluminescence (ECL) detection system.

#### Isothermal titration calorimetry

Isothermal titration calorimetry (ITC) experiments were performed using the PEAQ-ITC system (Malvern) in 20 mM HEPES/NaOH pH 7.5 and 150 mM NaCl at 10 °C. Mic60 concentration in the sample cell varied between the variants in the range of 44-81 µM, Mic19 concentration in the syringe between 391-809 µM. The Malvern analysis software was used to integrate the binding isotherms and calculate the binding parameters.

#### Liposome co-sedimentation assay

Folch lipids (total bovine brain lipids fraction I, Sigma) were dried under an argon stream, dissolved in 20 mM HEPES/NaOH pH 7.5 and 150 mM NaCl, incubated overnight at 4 °C and sonified for 15 min in a sonification bath. 40 µl of a reaction mixture containing 0.6 mg/ml liposomes and 5 µM protein (or complex) were incubated for 30 min at room temperature (RT) and centrifuged at 200,000 *g* for 16 min at 20 °C. The respective supernatant and pellet fractions were analyzed by SDS-PAGE and the protein bands quantified using ImageJ (version 1.50i, (*66*)).

#### Analytical ultracentrifugation

The fusion constructs Mito2_CHCH and Mito2_CHCH^L676D^ were analyzed at protein concentrations of 0.05 – 1 mg/ml in 20 mM HEPES/NaOH pH 7.5 and 150 mM NaCl using a Beckman Optima XL-I centrifuge equipped with an An50Ti rotor and double sector cells. Sedimentation equilibrium measurements were carried out at 20 °C and 16,000 rpm. The data were recorded at a wavelength of 230 or 280 nm and analyzed using the software Sedfit (*67*). No concentration-dependent assembly was observed for either construct in the applied concentration range.

#### *S. cerevisiae* strains and plasmids

*Saccharomyces cerevisiae* strains are derivates of YPH499 (*68*). Deletion strains *mic60Δ* and *mic19Δ* were described previously (*23*). For generation of different Mic60 and Mic19 variants containing individual amino acid substitutions, PCR fragments containing either the *MIC60* or the *MIC19* open reading frames together with their natural promoter and terminator regions were cloned into plasmid pRS416 and the respective mutations were generated via site-directed mutagenesis (see Extended Data Table S3 for a list of plasmids and Extended Data Table S4 for a list of *S. cerevisiae* strains used in this study). *S. cerevisiae* strains expressing Mic19 fused to a C-terminal Protein A tag for immunoprecipitation were generated by homologous recombination using a transformation cassette that consists of a tobacco etch virus protease cleavage site, a ZZ domain of *S. aureus* Protein A for immunoglobulin G (IgG) binding, and a *HIS3* marker gene for selection (*69*). For isolation of mitochondria, *S. cerevisiae* cells were grown in liquid minimal glycerol medium (0.67% [w/v] yeast nitrogen base, 0.07% [w/v] CSM amino acid mix minus uracil, 3% [v/v] glycerol) at 30 °C.

#### Isolation of *S. cerevisiae* mitochondria

Cells were grown in minimal glycerol medium to mid-log phase and harvested by centrifugation at 1,200 *g* for 5 min at RT. Pellets were resuspended in 2 ml/g wet weight DTT softening buffer (0.1 M Tris/H2SO4 pH 9.4; 10 mM DTT) and incubated for 20 min at 30 °C. After centrifugation (2,000 *g* for 5 min at RT), cell pellets were washed with Zymolyase buffer (1.2 M sorbitol; 20 mM KPi pH 7.4). Cells were resuspended in 6.5 ml Zymolyase buffer containing 4 mg of Zymolyase per gram of cells (wet weight) and incubated for 30 min at 30 °C for enzymatic digestion of the cell wall. The resulting spheroplasts were harvested by centrifugation (2,000 *g* for 5 min at RT), washed again with Zymolyase buffer and resuspended in 6.5 ml of homogenization buffer (0.6 M sorbitol; 10 mM Tris/HCl pH 7.4; 1 mM EDTA; 0.2% [w/v] bovine serum albumin [BSA]; 1 mM phenylmethylsulfonyl fluoride [PMSF]) per gram of cells. Spheroplasts were then homogenized using a glass-teflon Dounce homogenizer (15 strokes). The suspension was centrifuged (1,500 *g* for 5 min at 4 °C) to remove cell debris. The supernatant was transferred into a new tube and again centrifuged (15,000 *g* for 10 min at 4 °C). The mitochondria-containing pellet was resuspended in SEM buffer (250 mM sucrose; 10 mM MOPS pH 7.2; 1 mM EDTA). Protein concentration was measured via a Bradford assay and adjusted to 10 mg total protein / ml suspension. Isolated mitochondria were flash frozen in liquid nitrogen and stored at -80 °C.

#### Steady state levels of mitochondrial proteins

For comparing the steady state levels of individual proteins in wild-type and mutant mitochondria, frozen samples were thawed slowly on ice and centrifuged (15,000 *g* for 10 min at 4 °C). The mitochondrial pellet was resuspended in Laemmli buffer (60 mM Tris/HCl pH 6.8; 2% [w/v] sodium dodecylsulfate [SDS]; 10% [v/v] glycerol; 0.01% bromophenole blue; 1% β-mercaptoethanol). The samples were incubated for 10 min at 65°C and loaded on SDS-PAGE gels for protein separation. Subsequently, proteins were blotted on polyvinylidene difluoride (PVDF) membranes and visualized using specific antibodies together with an enhanced chemiluminescence (ECL) detection system (see Extended Data Table S5 for a list of the used antibodies).

#### Immunoprecipitation

For IgG affinity chromatography, 0.9 mg of isolated *S. cerevisiae* mitochondria (total protein content) containing Protein A-tagged Mic19 were resuspended in solubilization buffer (20 mM Tris/HCl pH 7.4; 50 mM NaCl; 0.1 mM EDTA; 10% [v/v] glycerol; 2 mM PMSF, 1x Roche protein inhibitor cocktail; 1% [w/v] digitonin) and incubated for 30 min at 4°C. Mitochondrial detergent extracts were centrifuged (20,000 *g* for 10 min at 4 °C), and the supernatant was incubated with 50 µl human IgG-coupled Sepharose beads (pre-equilibrated with 0.5 M acetate, pH 3.4) for 90 min at 4 °C. Beads were washed 10x with digitonin buffer (20 mM Tris/HCl pH 7.4, 0.5 mM EDTA, 60 mM NaCl, 10% [v/v] glycerol, 2 mM PMSF, 0.3% digitonin) followed by centrifugation (700 *g* for 30 s at 4 °C). Bound proteins were eluted by TEV protease cleavage over night at 4°C and centrifugation (1,200 g for 30 s at 4 °C).

#### Electron microscopy of *S. cerevisiae* mitochondria

*S. cerevisiae* cells grown in minimal medium supplemented with 2% glucose for 24 h at 30 °C were diluted in minimal glycerol medium and grown until early log phase. Cells were handled as previously described (*70*). Cells were fixed for 3 h with freshly prepared 4% [w/v] paraformaldehyde and 0.5% (v/v) GA in 0.1 M citrate buffer adjusted to growth conditions for pH and temperature (71). After washing with citrate buffer, cells were permeabilized for 1 h with 1% [w/v] sodium metaperiodate at 4 °C. Cells were washed with 0.1 M phosphate buffer and then embedded in 12% gelatine by cooling the 37 °C warm gelatine in ice after 10 min of incubation. 1 mm^3^ cubes were cut, infiltrated overnight with 2.3 M sucrose, mounted onto specimen chucks and frozen in liquid nitrogen. Ultrathin sections were cut using an UC7 ultramicrotome (Leica) at -110 °C and collected on formvar/carbon-coated copper grids (Plano). The gelatine was removed by washing with phosphate buffered saline at 37 °C and water prior to staining with 3% [w/v] silicotungstic acid hydrate (Fluka) in 2.8% [w/v] polyvinyl alcohol (Sigma-Aldrich) in water for 5 min. Grids were imaged after drying with the transmission electron microscope EM910 (Zeiss) operating at 80 kV and equipped with a Quemesa CCD camera and the imaging software iTEM (Emsis) at 10,000 × magnification. All data are presented as the mean ± the s. d. and value differences were compared statistically. Data analysis and plotting was done with the statistic program *R*. The normal distribution was tested using the Kolmogorov-Smirnov test as well as the Q-Q plot. Since data was not normally distributed, the two-sided Wilcoxon-Rank-sum test for independent samples with continuity correction was used. For all groups, n = 100 mitochondrial cross sections were analyzed. Differences of p ≤ 0.05 were considered significant (p ≤ 0.05*, p ≤ 0.01**, p ≤ 0.001***). *S. cerevisiae* cells used for tomography analysis were harvested at 2,000 rpm using a Stat Spin Microprep 2 table top centrifuge. After centrifugation, the pellet was fixed by immersion using 2% glutaraldehyde in 0.1 M cacodylate buffer at pH 7.4. Fixation was performed for 60 min at RT. The fixed pellet was immobilized with 2% agarose in sodium cacodylate buffer at pH 7.4. Small pieces of the immobilized pellet were fixed using buffered 1% osmium tetroxide followed by aqueous 1% uranyl acetate, samples were dehydrated and embedded in Agar 100. Alternatively, pelleted cells were vitrified using a BAL-TEC HPM-010 high-pressure freezer. The samples were substituted over 72 h at -90 °C in a solution of 2% OsO4, 0.1% uranyl acetate and 5% H_2_O in anhydrous acetone. After a further incubation over 20 h at -20°C, samples were warmed up to +4 °C and washed with anhydrous acetone subsequently. The samples were embedded at RT in Agar 100 (Epon 812 equivalent) at 60 °C over 24 h. After ultrathin sectioning (230 nm), section were counterstained with lead citrate. Images were taken in a Talos L120C transmission electron microscope (Thermo Fischer Eindhoven, The Netherlands). Tilt series from 230 nm thick sections were recorded in 4K mode using the dose symmetric scheme from -65° to 65° at 2° intervals. Tomograms were calculated using Etomo (http://bio3d.colorado.edu/) (*72*). Size determination of CJs were calculated using 3dmod.

#### Preparation of the Mic60-Mic19 model within crista junctions

The model of Mic60 and Mic19 in crista junctions was created and prepared by Dr. Erik Werner (2021), RNS Berlin (www.rns.berlin). The model comprises the crystal structures of the coiled-coil region of Mic60 from *L. thermotolerans* (ltMic60CC, 235-382), the LBS 1 and mitofilin domain of Mic60 from *Chaetomium thermophilum* and the CHCH domain of Mic19 from *C. thermophilum* (Mito2_CHCH). In this, the sequence of *Lachancea thermotolerans* Mic60 was used as template. All other parts of Mic60 were modelled as unstructured regions. A crista junction diameter of 17.5 nm was used (*20*). The model was generated using the Maya® software from Autodesk, Inc. (https://www.autodesk.com/products/maya/) and Modeling kit and Rigging kit of the plugin Molecular Maya (mMaya) from Digizyme, Inc. (https://clarafi.com/tools/mmaya/).

**Fig. S1.**
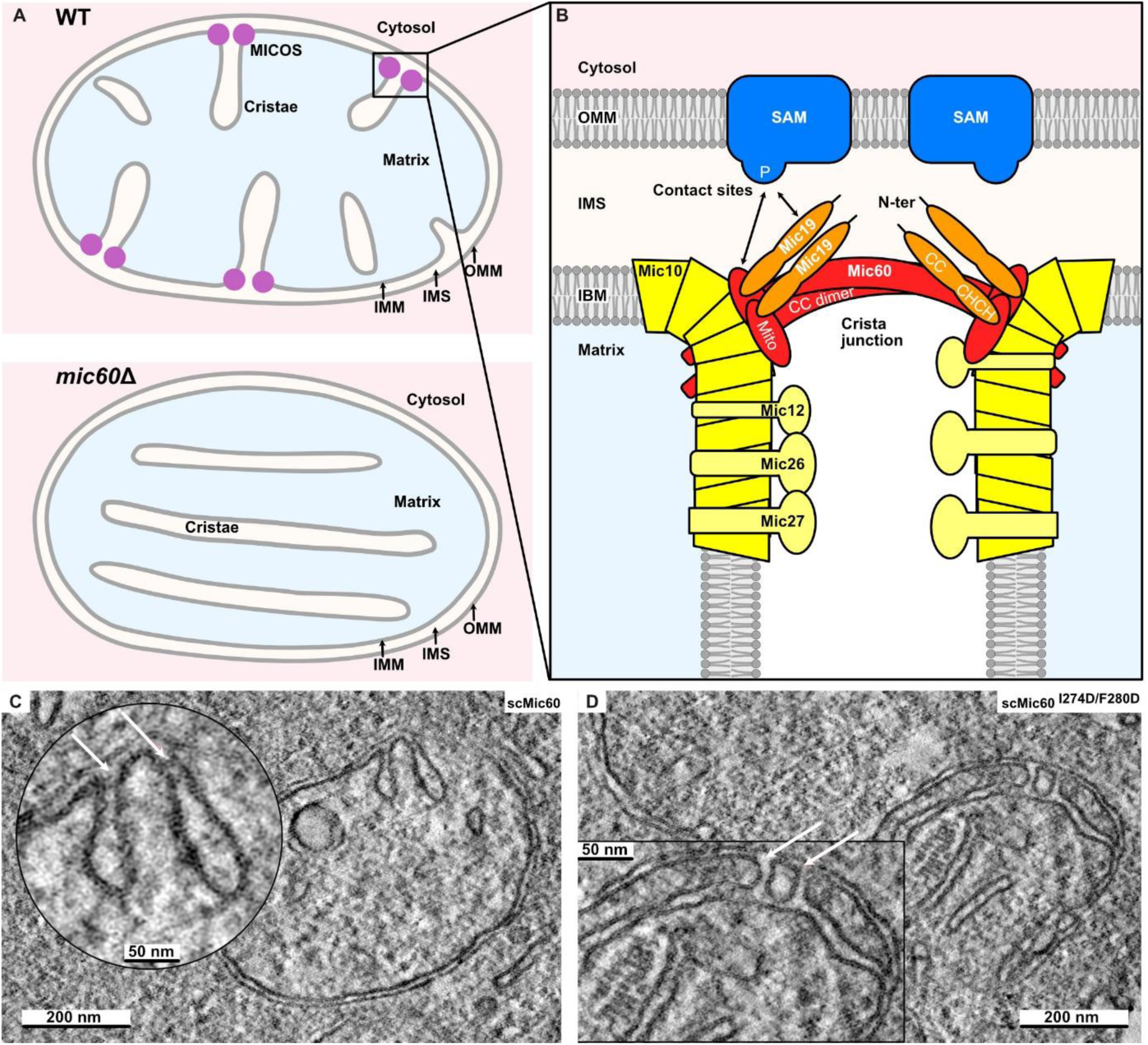
Overview of mitochondrial architecture. **(A)** Mitochondria are surrounded by two membrane systems. The outer mitochondrial membrane (OMM) envelopes the organelles and mediates communication and exchange of molecules with the cytosol and other membrane-bound intracellular compartments. The inner mitochondrial membrane (IMM) consists of two structurally and functionally distinct domains. The inner boundary membrane (IBM) is in close proximity to the OMM, comprising the aqueous intermembrane space (IMS) compartment. Large membrane areas, termed cristae, extend from the IMM and wrinkle into the matrix. These cristae membranes adopt the shape of branched tubules, sheets or discs and determine the characteristic ultrastructure of mitochondria. IBM and cristae membranes largely differ in their protein content. Whereas the IBM particularly contains metabolite carriers and translocation machineries for macromolecules, like polypeptides, cristae membranes are exceptionally protein-rich and harbor the respiratory chain complexes and F1Fo-ATP synthase for oxidative phosphorylation. The connections between cristae and IBM are termed crista junctions (CJs). These specialized membrane regions exhibits an exceedingly high curvature and are thought to act as diffusion barriers for metabolites and proteins. **(B)** The formation and stabilization of crista junctions is mediated by an evolutionary conserved hetero-oligomeric protein complex, the mitochondrial contact site and cristae organizing system (MICOS). The core complex consists of six subunits in yeast and seven in metazoa. MICOS is made of a Mic60 and a Mic10 module with distinct properties and functions. Mic60 (formerly known as mitofilin, IMMT or Fcj1) is the centerpiece of the membrane-bridging subcomplex that forms contact sites between IMM and OMM through interactions with partner protein complexes, like the sorting and assembly machinery (SAM). The Mic60 protein consists of an N-terminal transmembrane segment anchored in the IMM and a large hydrophilic domain in the IMS that exhibits an extended coiled-coil region and a C-terminal mitofilin signature domain. Mic60 is firmly associated with Mic19, a peripheral membrane protein that may act as a redox sensor through an intramolecular disulfide bond. In metazoa, the Mic60 module additionally contains Mic25 that belongs to the same protein family as Mic19. Studies in yeast and human mitochondria suggest that the N-terminal domain of Mic19 and the parts of the Mic60 IMS domain differentially contribute to membrane contact site formation together with the polypeptide transport-associated (POTRA) domain of Sam50. Homo-oligomers of Mic10 form the backbone structure of the second MICOS subcomplex and contribute to the formation and stabilization of membrane curvature at crista junctions. The oligomeric state of Mic10 is regulated by Mic26 and Mic27 in an antagonistic manner and modulated by the phospholipid cardiolipin. The two MICOS modules are connected by Mic12 in yeast or Mic13/QIL1 in metazoa. P: POTRA domain of Sam50, N-ter. Mic19: N-terminal region of Mic19, CC: coiled-coil domain, CHCH: coiled-coil-helix-coiled-coil-helix domain, Mito: mitofilin domain, CC dimer: Mic60 coiled-coil domain dimer. Ablation of MICOS leads to the collapse of crista junctions and the detachment of cristae from the IBM **(A**, **bottom)**. Cristae accumulate as lamellar membrane stacks in the matrix. MICOS-deficient mitochondria show defects in oxidative phosphorylation and a variety of stress responses. **(C)** EM micrograph of mitochondria from fixed *mic60Δ S. cerevisiae* reconstituted with Mic60. Filamentous density near the CJ is observed in the majority of analyzed CJs (45 of 55 CJs, white arrows). **(D)** Same as in **c**, but reconstituted with the tetramerization mutant Mic60^I274D/F280D^. Interference with Mic60 tetramerization results in a reduction of CJs. From the remaining CJs (white arrows), only 4 out of 29 showed a filamentous density. The inserts show magnifications.

**Fig. S2.**
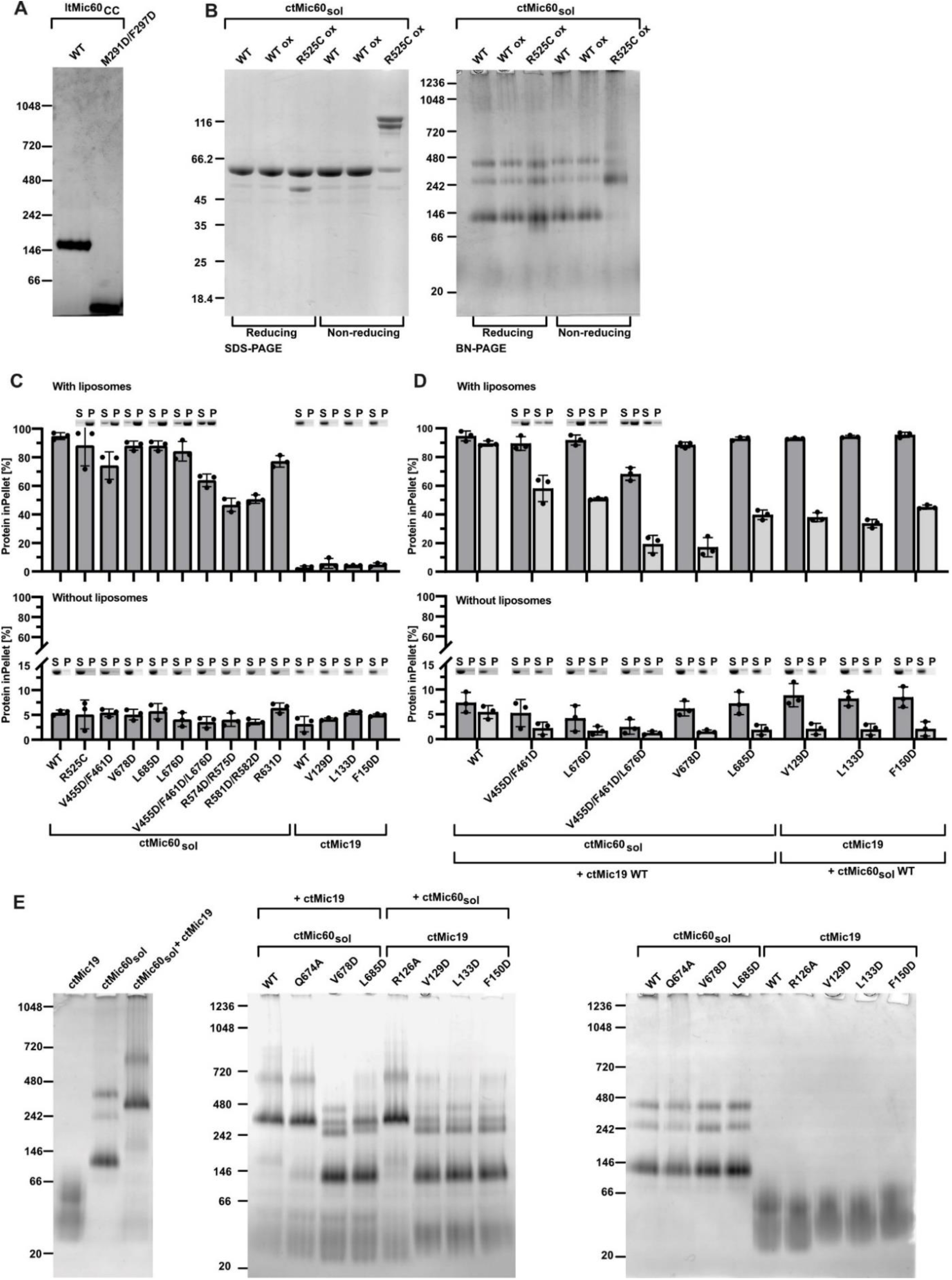
Oligomerization and interaction of Mic60 and Mic19. **(A)** BN-PAGE of purified ltMic60CC and the tetramerization mutant M291D/F297D. **(B)** SDS-PAGE (left) and BN-PAGE (right) analysis of the CtMic60_sol_ and CtMic60_sol_^R525C^ under reducing and non-reducing conditions. WT represents CtMic60_sol_ under non-oxidized conditions, whereas WT Ox and R525C Ox show CtMic60_sol_ and the R525C mutant after oxidation using CuSO4. **(C), (D)** SDS-PAGE analysis and quantification of liposome co-sedimentation assay of different CtMic60_sol_/ctMic19 single variants **(C)** and respective complexes **(D)**. S, supernatant; P, pellet. The light grey bars in **(D)** represent ctMic19 and its variants and the dark grey bars CtMic60_sol_ and its variants. Measurements were done in triplicate and error bars indicate the s.d. of each data set. SDS-PAGE data shown in Fig. 3c and 4e are not included here. **(E)** BN-PAGE analysis CtMic60_sol_ variants and their complexes with ctMic19 variants. Proteins and protein complexes in all figures were visualized with Coomassie Brilliant Blue.

**Fig. S3.**
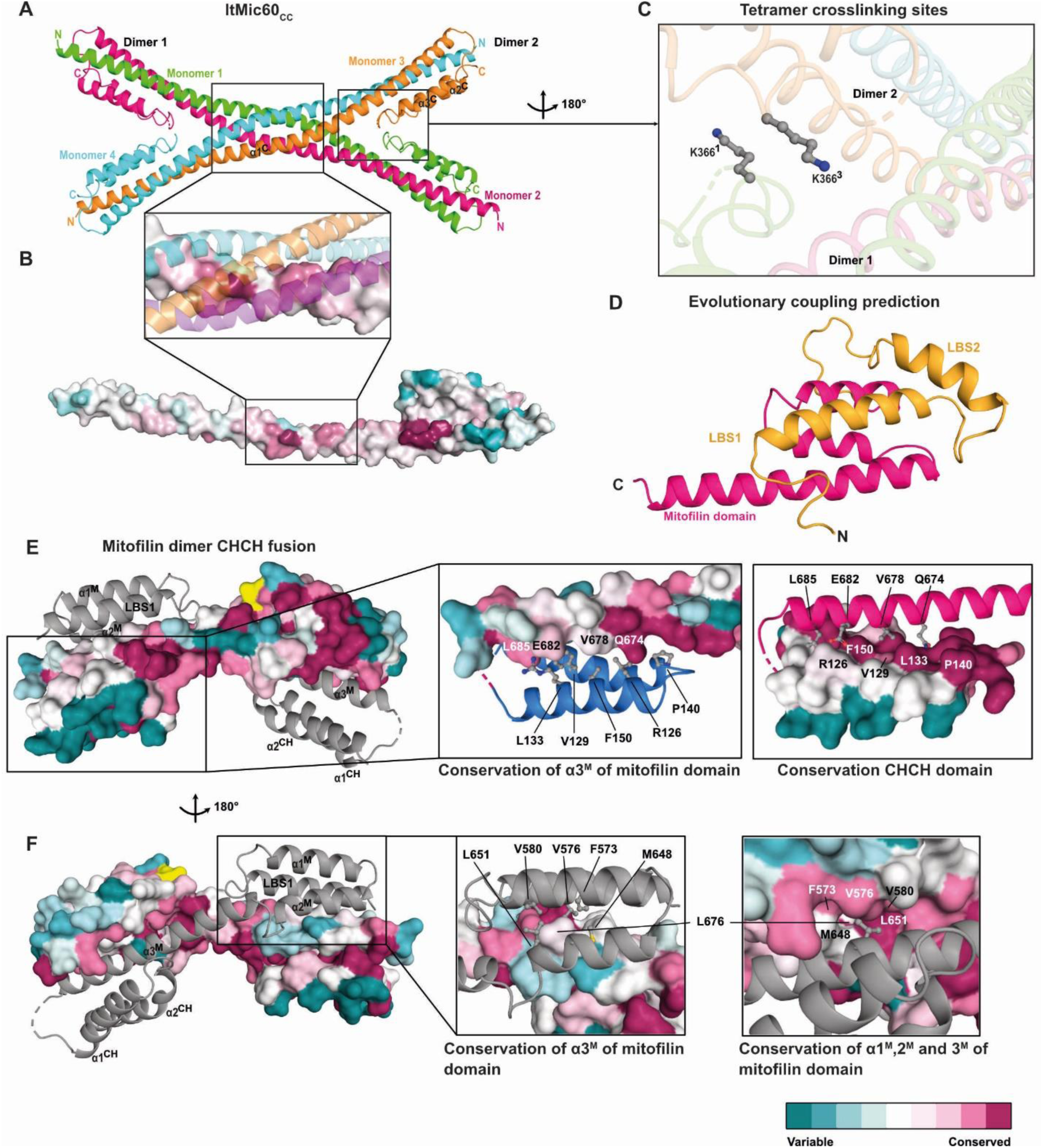
Structural details of Mic60 and Mic19. **(A)** Tetrameric structure of ltMic60CC. Tetrameric helical assemblies are often found in membrane-remodeling proteins, such as in SNARE complexes^46^ or the stalks of dynamin proteins^47^, although the detailed topologies differ. **(B)** Surface conservation plot of the tetramer interface (zoom) and the monomer of ltMic60CC. The other monomers are shown in cartoon representation. Conserved residues are colored dark magenta, variable residues dark cyan. **(C)** Localization of K366, which was exchanged to cysteine to stabilize the tetramer via a disulfide bridge. K366^1^ refers to monomer 1 and K366^3^ to monomer 3. **(D)** Structure prediction of the C-terminal region of Mic60 from co-evolution analysis. The mitofilin domain is colored in magenta and the LBS 1 and LBS 2 in orange. **(E), (F)** Surface conservation plot of one monomer of Mito2_CHCH. The second monomer is shown in cartoon representation. The GS-linker is colored in yellow. Both magnifications in **(E)** show the conservation of the mitofilin-CHCH domain interface. The CHCH domain or helix α3^M^ from the mitofilin domain are shown as cartoon representation for better visualization. The magnifications in **(F)** represent the conservation of the mitofilin-dimer interface.

**Fig. S4.**
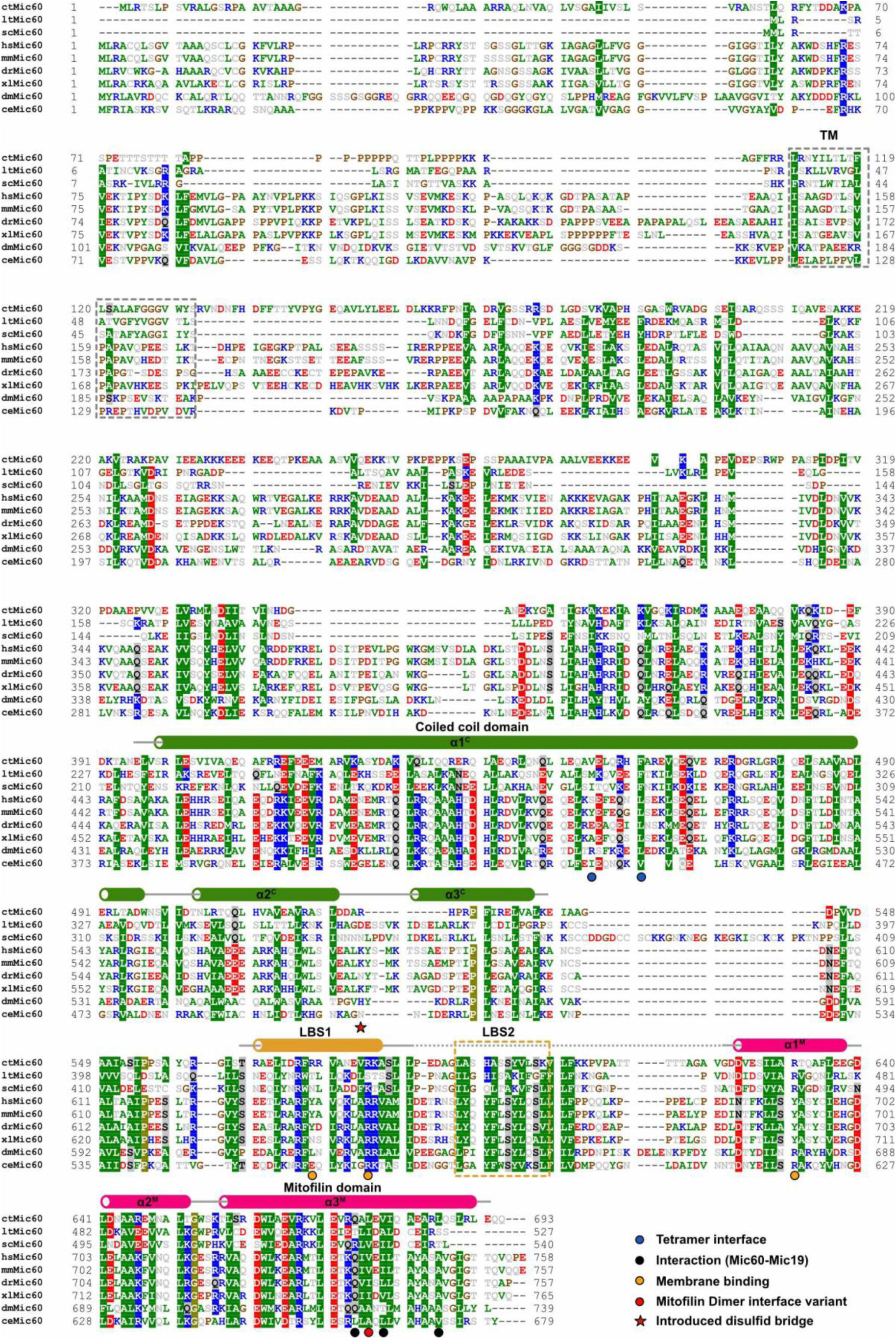
Sequence alignment of Mic60. The following sequences have been aligned: *Chaetomium thermophilum* (ctMic60, Uniprot accession code G0SHY5), *Lachancea thermotolerans* (ltMic60, C5E325), *Saccharomyces cerevisiae* (scMic60, P36112), *Homo sapiens* (hsMic60, Q16891), *Mus musculus* (mmMic60, Q8CAQ8), D*anio rerio* (drMic60, Q6PFS4), *Xenopus laevis* (xlMic60, A0A1L8HKP3), *Drosophila melanogaster* (dmMic60, P91928) *Caenorhabditis elegans* (ceMic60, Q22505). Amino acids are colored according to their chemical and physical properties (positive charge: blue, negative charge: red, hydrophobic: green, proline and glycine: brown, all others: grey). Conservation of more than 70% of all sequences is indicated by the highlighted background. Residues involved in tetramerization are labelled with 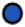, in interaction with Mic19 with 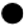, in membrane binding with 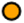, in mitofilin dimerization with 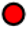 and the residue chosen for cysteine replacement with 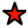.

**Fig. S5.**
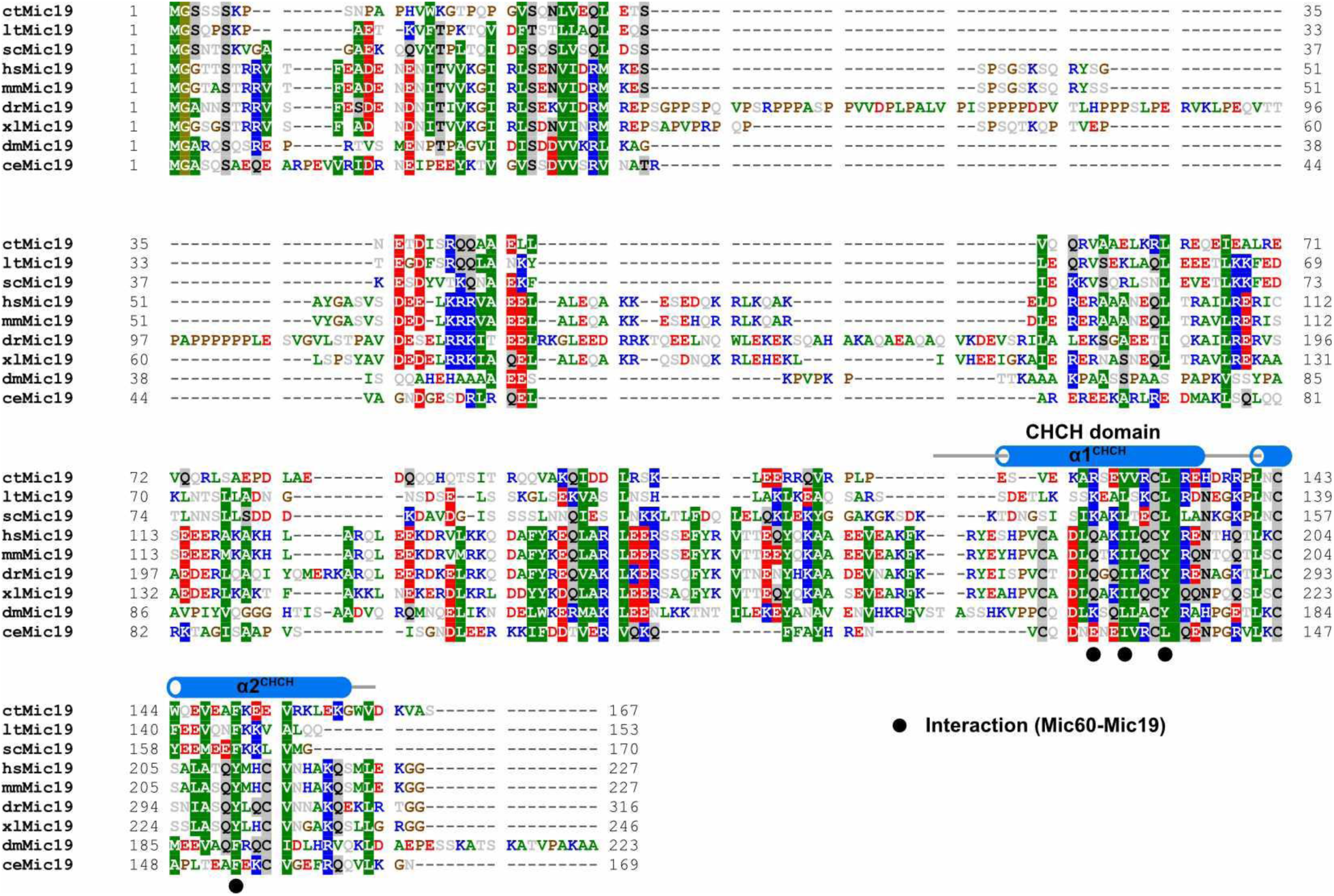
Sequence alignment of Mic19. The following sequences have been aligned: *Chaetomium thermophilum* (ctMic19, Uniprot accession code G0S140), *Lachancea thermotolerans* (ltMic19, C5E3G4), *Saccharomyces cerevisiae* (scMic19, P43594), *Homo sapiens* (hsMic19, Q9NX63), *Mus musculus* (mmMic19, Q9CRB9), *Danio rerio* (drMic19, Q502T3), *Xenopus laevis* (xlMic19, Q7ZYP1), *Drosophila melanogaster* (dmMic19, Q9VA18), *Caenorhabditis elegans* (ceMic19, Q21551). Amino acids are colored according to their chemical and physical properties (positive charge: blue, negative charge: red, hydrophobic: green, proline and glycine: brown, all others: grey). Conservation of more than 70% of all sequences is indicated by the highlighted background. Residues involved in interaction with Mic60 are labelled with 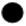.

**Fig. S6.**
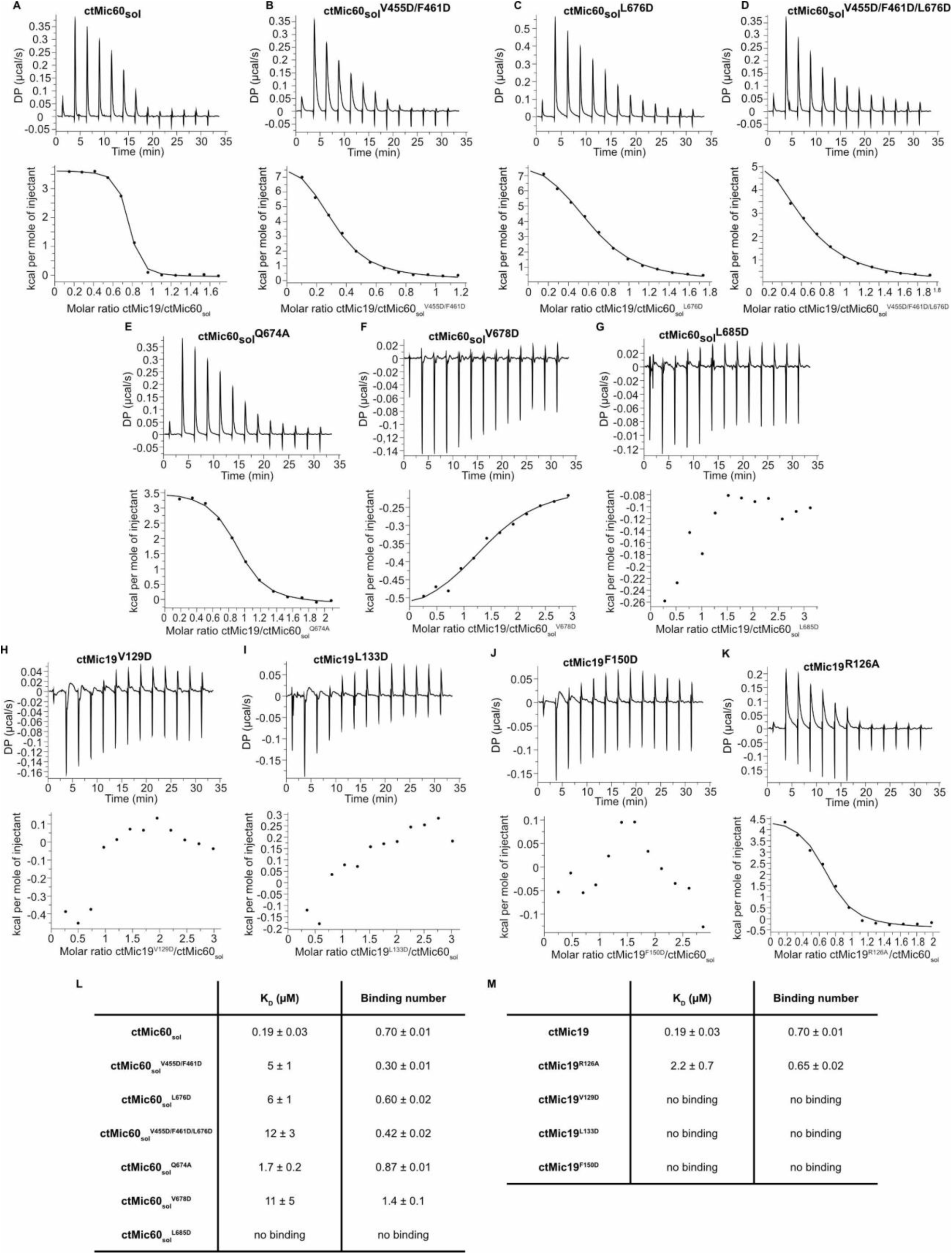
ITC data of Mic60 and Mic19. (**A-K**) ITC experiments of different ctMic60_sol_ variants and ctMic19 variants. At 10 °C, a concentrated solution of Mic19 in the syringe was titrated to Mic60 present in the sample cell, and the resulting heat change was monitored. Mic60 concentration in the sample cell varied between 44-81 µM and Mic19 concentrations between 390-810 µM. **(L-M)** Overview of K_D_s (µM) and binding numbers of the ITC experiments shown in **(A-K)**. The deviation represents the root-mean-square error of the fit (*n*=1). **(L)** includes the ctMic60_sol_ variants, which have been titrated with ctMic19 and (M) the ctMic19 variants, which have been titrated into ctMic60_sol_.

**Fig. S7.**
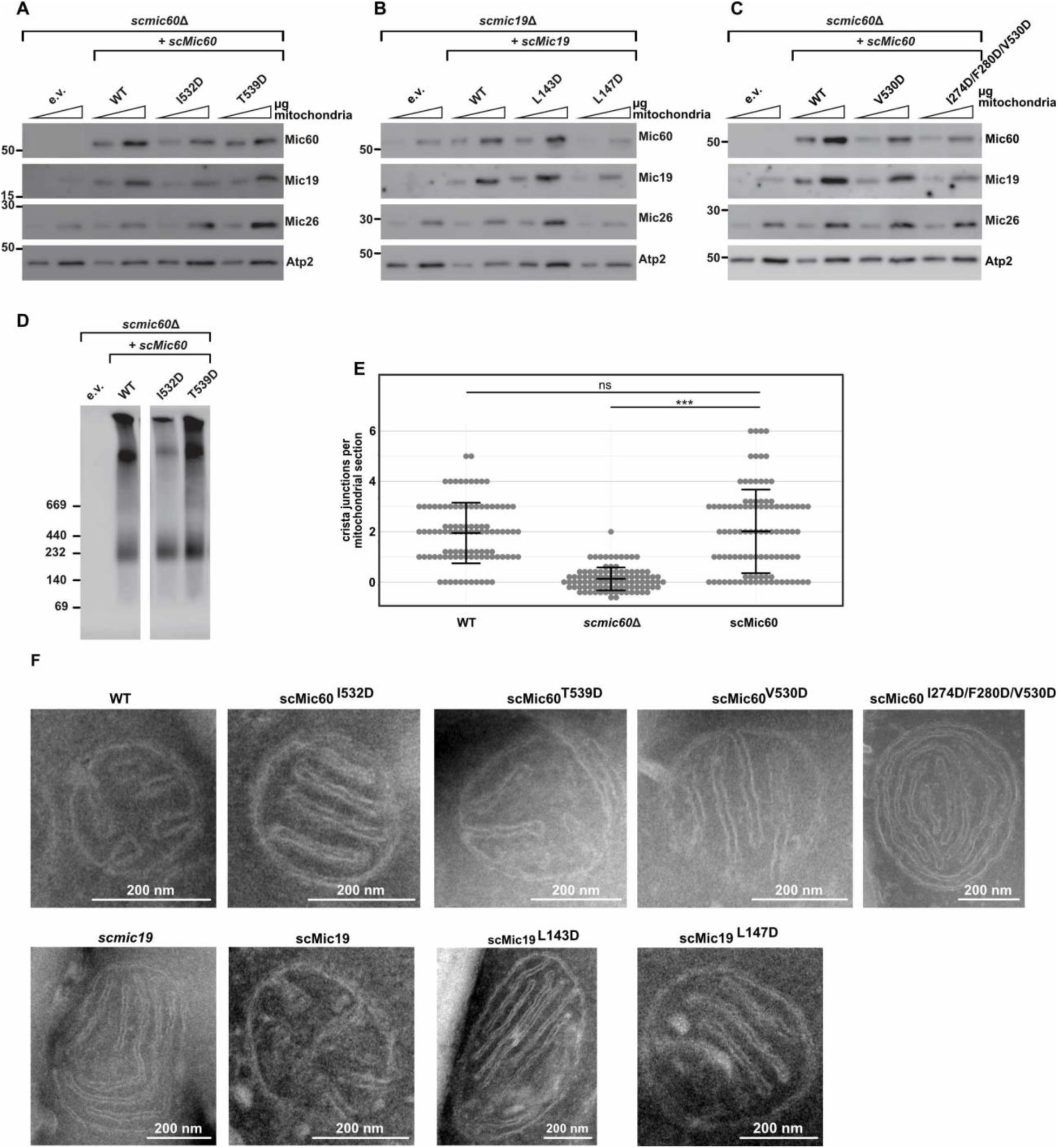
***S. cerevisiae* in vivo analysis. (A)** Comparison of steady state protein levels in isolated mitochondria of *S. cerevisiae* strains lacking the chromosomal *MIC60* gene (*mic60*Δ) transformed with an empty vector (e.v.), or plasmids encoding either wild-type (WT) scMic60 or the scMic60 variants I532D and T539D, respectively. Mitochondrial samples were subjected to SDS-PAGE and immunoblotting with the indicated antisera. Note that the I532D variant was expressed at lower levels compared to WT. **(B)** Protein steady state level analysis as in **(A)** using mitochondria of *S. cerevisiae* strains lacking Mic19 (*mic19*Δ) or expressing WT scMic19 or the scMic19 L143D and L147D variants, respectively. Note that the L147D variant was expressed at lower levels compared to WT. Accordingly, also lower levels of the MICOS components Mic60 and Mic26 were observed. **(C)** Experiment as described in **(A)** analyzed scMic60 variants were V530D and I274D/F280D/V530D. The Mic60 variant with three amino acid substitutions was expressed at slightly lower levels compared to WT. **(D)** Mitochondria as described in **(A)** additionally containing a Protein A-tagged variant of Mic19, were solubilized in digitonin-containing buffer and subjected to IgG affinity chromatography, Elution fractions were analyzed by blue native PAGE und immunodetection with antibodies raised against scMic60. **(E)** Number of crista junctions per mitochondrial section. N = 100 mitochondrial cross sections were counted with maximally two mitochondria from the same cell. Each data point represents one mitochondrial cross section, and the mean and the standard deviation are shown. Statistically significant differences in the mean values are indicated. **(F)** Representative electron micrographs of *S. cerevisiae* mitochondria in ultrathin sections of cells expressing the indicated scMic60 and scMic19 variants. scMic19 represents the *mic19*Δ strain complemented with plasmid pRS416-scMic19 (WT) (see also Fig. 2e-g).

---

**Table S1.**
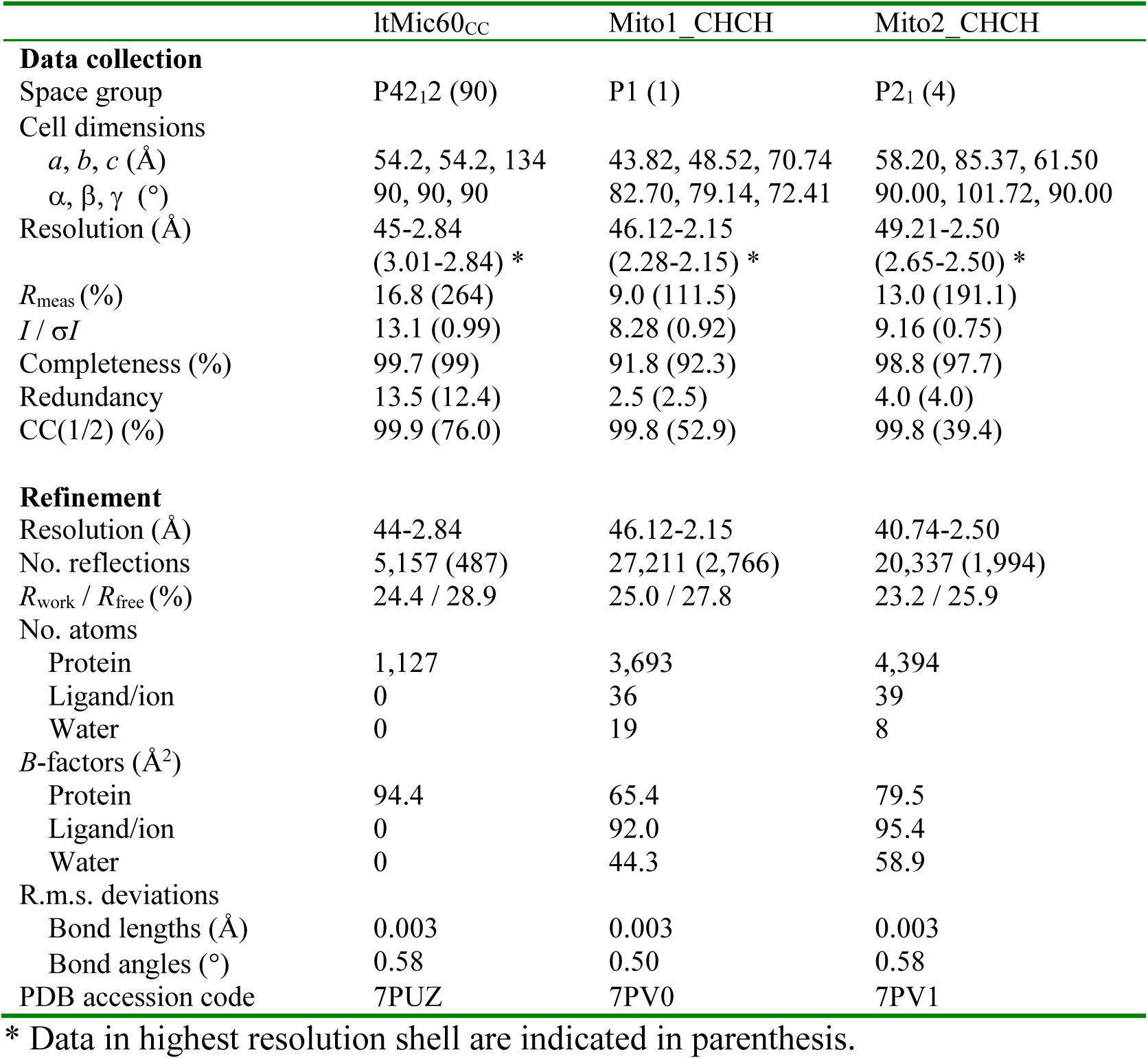
Crystallographic data collection and refinement statistics.

**Table S2.** Overview of Mic60 and Mic19 constructs. Summary of all constructs used in this study. Lt: *Lachancea thermotolerans*, ct: *Chaetomium thermophilum*, sc: *Saccharomyces cerevisiae*

| ltMic60 |  | ctMic60 |  |  | scMic60 |  |  | Comments |
| --- | --- | --- | --- | --- | --- | --- | --- | --- |
| Boundaries | Construct name | Boundaries | Mutations | Construct name | Boundaries | Mutations | Construct name |  |
| 207-382 | ltMic60cc |  |  |  |  |  |  | Crystallized coiled-coil domain |
|  |  | 208-691 |  | ctMic60<sub>sol</sub> | 1-540 |  | scMic60 | Wild type protein |
|  |  | 208-691 | V455D/F461D | ctMic60<sub>sol</sub><sup>V455D/F461D</sup> | 1-540 | I274D/F280D | scMic60<sup>I274D/F280D</sup> | Broken tetramer interface |
|  |  | 208-691 | L676D | ctMic60<sub>sol</sub><sup>L676D</sup> | 1-540 | V530D | scMic60<sup>V530D</sup> | Broken mitofilin dimer interface |
|  |  | 208-691 | V455D/F461D/L676D | ctMic60<sub>sol</sub><sup>V455D/F461D/L676D</sup> | 1-540 | I274D/F280D/V530D | scMic60<sup>I274D/F280D/V530D</sup> | Broken tetramer and mitofilin dimer interface |
|  |  | 208-691 | Q674A | ctMic60<sub>sol</sub><sup>Q674A</sup> |  |  |  | Disturbed Mic60-Mic19 interaction |
|  |  | 208-691 | V678D | ctMic60<sub>sol</sub><sup>V678D</sup> | 1-540 | I532D | scMic60<sup>I532D</sup> | Disturbed Mic60-Mic19 interaction |
|  |  | 208-691 | L685D | ctMic60<sub>sol</sub><sup>L685D</sup> | 1-540 | T539D | scMic60<sup>T539D</sup> | Disturbed Mic60-Mic19 interaction |
|  |  | 208-691 | R574D/R575D | ctMic60<sub>sol</sub><sup>R574D/R575D</sup> |  |  |  | Reduced membrane binding |
|  |  | 208-691 | R581D/K582D | ctMic60<sub>sol</sub><sup>R581D/K582D</sup> |  |  |  | Reduced membrane binding |
|  |  | 208-691 | R631D | ctMic60<sub>sol</sub><sup>R631D</sup> |  |  |  | Reduced membrane binding |
|  |  | ctMic19 |  |  | scMic19 |  |  | Comments |
|  |  | Boundaries | Mutations | Construct name | Boundaries | Mutations | Construct name |  |
|  |  | 1-164 |  | ctMic19 | 1-170 |  | scMic19 | Wild type protein |
|  |  | 1-164 | R126A | ctMic19<sup>R126A</sup> |  |  |  | Disturbed Mic60-Mic19 interaction |
|  |  | 1-164 | V129D | ctMic19<sup>V129D</sup> | 1-170 | L143D | scMic19<sup>L143D</sup> | Disturbed Mic60-Mic19 interaction |
|  |  | 1-164 | L133D | ctMic19<sup>L133D</sup> | 1-170 | L147D | scMic19<sup>L147D</sup> | Disturbed Mic60-Mic19 interaction |
|  |  | 1-164 | F150D | ctMic19<sup>F150D</sup> |  |  |  | Disturbed Mic60-Mic19 interaction |
| Fusion proteins |  | ctMic60 | ctMic19 | Mutation | Fusion protein |  |  | Comments |
| Mito1_CHCH |  | 624-691 | 116-164 |  | ctMic60<sup>624-691</sup>-GSGS-ctMic19<sup>116-164</sup> |  |  | Crystallized mitofilin and CHCH domain |
| Mito2_CHCH |  | 565-586-GS-622-691 | 116-164 |  | ctMic60<sup>565-586-GS-622-691</sup>-GSGS-ctMic19<sup>116-164</sup> |  |  | Crystallized mitofilin-dimer and CHCH domain |
| Mito2_CHCH<sup>L676D</sup> |  | 565-586-GS-622-691 | 116-164 | L676D | ctMic60<sup>565-586-GS-622-691</sup>-GSGS-ctMic19<sup>116-164</sup> |  |  | Broken mitofilin domain dimer and CHCH domain |

**Table S3.** Plasmids used in this study.

| **Plasmid** | **Description** | **Reference** |
| --- | --- | --- |
| pRS416 | CEN, empty vector |  |
| pRS416-Mic60 | CEN, Mic60 | This study |
| pRS416-Mic60<sup>I274D/F280D</sup> | CEN, Mic60<sup>I274D/F280D</sup> | This study |
| pRS416-Mic60<sup>V530D</sup> | CEN, Mic60<sup>V530D</sup> | This study |
| pRS416-Mic60<sup>I274D/F280D/V530D</sup> | CEN, Mic60<sup>I274D/F280D/V530D</sup> | This study |
| pRS416-Mic60<sup>I532D</sup> | CEN, Mic60<sup>I532D</sup> | This study |
| pRS416-Mic60<sup>T539D</sup> | CEN, Mic60<sup>T539D</sup> | This study |
| pRS416-Mic19 | CEN, Mic19 | This study |
| pRS416-Mic19<sup>L143D</sup> | CEN, Mic19<sup>L143D</sup> | This study |
| pRS416-Mic19<sup>L147D</sup> | CEN, Mic19<sup>L147D</sup> | This study |
| CEN - Centromer |  |  |

**Table S4.**
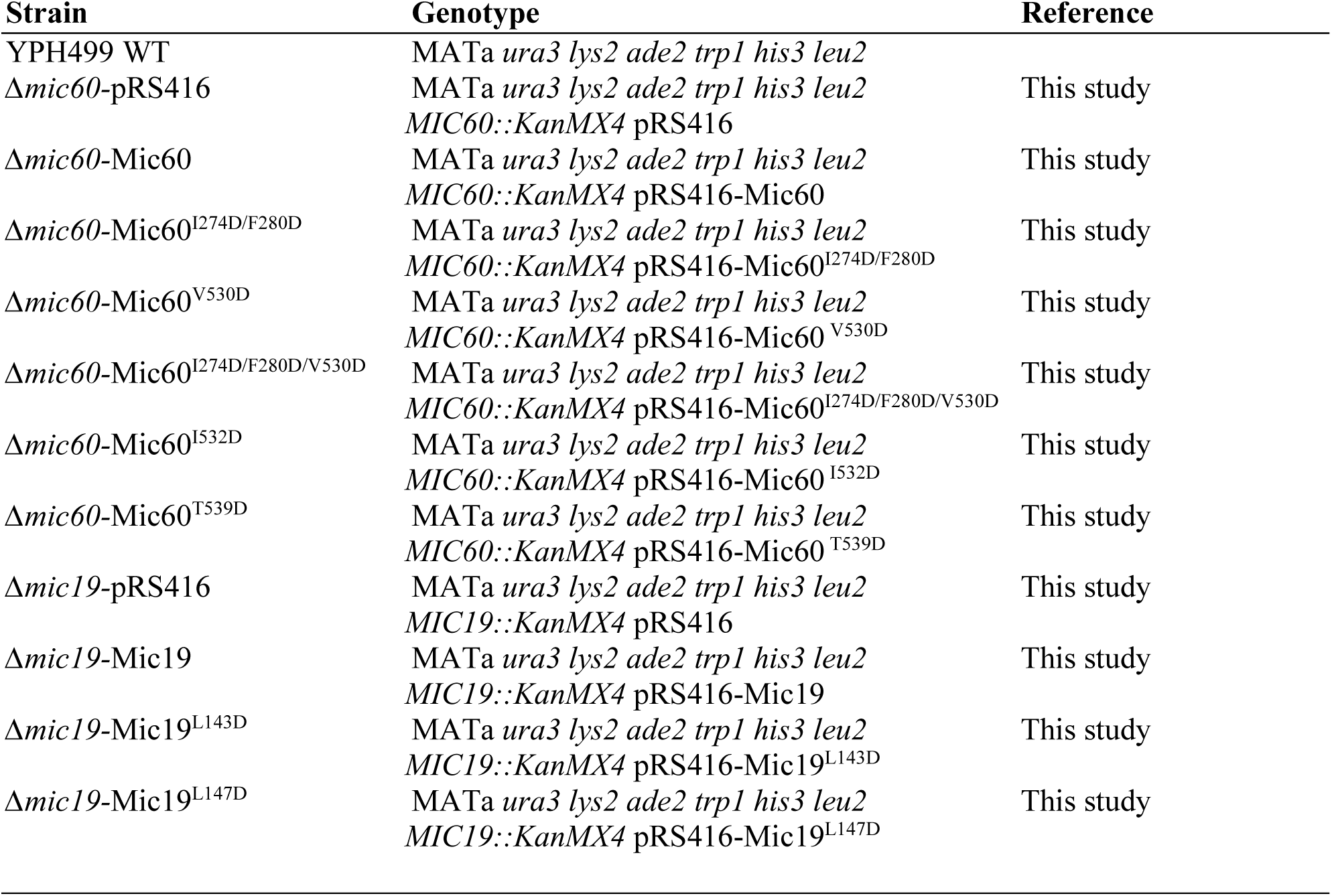
S. cerevisiae strains used in this study.

**Table S5.** Antibodies used in this study.

| **Antibodies** | **Source** | **Identifier** |
| --- | --- | --- |
| anti-Atp2 | Pfanner lab/ van der Laan lab |  |
| anti-Mic60 | This study; amino acids 140-295 | N/A |
| anti-Mic60 | This study; amino acids 168-295 | N/A |
| anti-Mic26 | Pfanner lab/ van der Laan lab | Ref. 23 |
| anti-Mic19 | Pfanner lab/ van der Laan lab | Ref. 23 |

